# Gut bacterial deamination of residual levodopa medication for Parkinson’s disease

**DOI:** 10.1101/2020.02.13.945717

**Authors:** Sebastiaan P. van Kessel, Hiltje R. de Jong, Simon L. Winkel, Sander S. van Leeuwen, Sieger A. Nelemans, Hjalmar Permentier, Ali Keshavarzian, Sahar El Aidy

## Abstract

Aromatic bacterial metabolites are attracting considerable attention due to their impact on gut homeostasis and host’s physiology. *Clostridium sporogenes* is a key contributor to the production of these bioactive metabolites in the human gut. Here, we show that *C. sporogenes* deaminates levodopa, the main treatment in Parkinson’s disease, and identify the aromatic aminotransferase responsible for the initiation of the deamination pathway. The deaminated metabolite from levodopa, 3-(3,4-dihydroxyphenyl)propionic acid, elicits an inhibitory effect on ileal motility in an *ex vivo* model. 3-(3,4-dihydroxyphenyl)propionic acid is detected in fecal samples of Parkinson’s disease patients on levodopa medication. Our data are of significant impact to the treatment of Parkinson’s disease, where constipation is reported as the most common gastrointestinal symptom. Overall, this study underpins the importance of the metabolic pathways of the gut microbiome involved in drug metabolism not only to preserve drug effectiveness, but also to avoid potential side effects of bacterial breakdown products of the unabsorbed residue of medication.

## Introduction

Gut bacteria produce a wide range of small bioactive molecules from different chemical classes, including aromatic amino acids (Donia and Fischbach, 2015). Bacterial products from aromatic amino acid degradation have been shown to play a critical role in intestinal barrier function, immune modulation and gut motility (Bansal et al., 2010; Bhattarai et al., 2018; Schiering et al., 2017; Venkatesh et al., 2014; Yano et al., 2015). In the lower part of the gastrointestinal (GI) tract, where oxygen is limited, aromatic amino acid degradation by anaerobic bacteria involves reductive or oxidative deamination (Barker, 1981) resulting in production of aromatic metabolites (Dodd et al., 2017; Elsden et al., 1976; Nierop Groot and De Bont, 1998; Yvon et al., 1997). Although the enzymes involved in the deamination pathway of the aromatic amino acids tryptophan, phenylalanine and tyrosine have been described (Dickert et al., 2000, 2002; Dodd et al., 2017), the enzyme involved in the initial transamination step remains unknown.

Recently, small intestinal (SI) microbiota have been implicated in the interference with levodopa drug availability (van Kessel et al., 2019; Maini Rekdal et al., 2019). Early *in vivo* studies showed that ~90% of levodopa is transported to the circulatory system (Bianchine et al., 1972; Morgan, 1971; Sasahara et al., 1981), leaving a ~10% unabsorbed fraction of residual levodopa that can act as substrate for other bacterial species associated with the lower, more anaerobic regions of the GI-tract (Goldin et al., 1973). Such bacterial-residual drug interaction might act as bioactive metabolites with an impact on gut homeostasis.

PD is often associated with non-motor symptoms especially in the GI-tract. GI-tract dysfunction such as constipation, drooling and swallowing disorders occur frequently in PD patients, especially constipation, which is reported in 80-90% of the PD patients (Fasano et al., 2015). Importantly, chronic idiopathic constipation is associated with SI motor abnormalities in the esophagus, stomach, jejunum and ileum (Panagamuwa et al., 1994; Van Der Sijp et al., 1993) and patients with constipation have a longer SI transit time compared to controls (Van Der Sijp et al., 1993). Only recently, SI dysfunction in PD was studied showing that the transit time in the SI was significantly longer in PD patients compared to healthy controls (HC) (Dutkiewicz et al., 2015; Knudsen et al., 2017). Using wireless electromagnetic capsules, the SI transit time was reported to be significantly higher in PD patients (400 min; n=22) compared to HC (295 min, n=15) (Knudsen et al., 2017).

This study uncovers the aminotransferase responsible for initiating the deamination pathway involved in the transamination of (among others) levodopa and shows that *C. sporogenes* can effectively deaminate levodopa to 3-(3,4-dihydroxyphenyl)propionic acid through the aromatic amino acid deamination pathway (Dodd et al., 2017). We show that the deamination product of gut bacterial degradation of the unabsorbed residues of levodopa in fecal samples from PD patients reduces ileal motility *ex vivo*. Our results highlight the urgency for further research on the effects of bacterial conversion of the unabsorbed residues of medication, which may affect host physiology.

## Results

### *Clostridium sporogenes* deaminates levodopa through its deamination pathway

*C. sporogenes* is able to deaminate proteinogenic aromatic amino acids (PAAA) through an anaerobic deamination pathway (**Figure 1A**) (Dickert et al., 2000, 2002; Dodd et al., 2017). We hypothesized that levodopa, a non-proteinogenic amino acid (NPAAA) and the main treatment in PD could be deaminated through the same pathway. Together with another NPAAA, 5-hydroxytryptophan (5-HTP, precursor of serotonin, over-the-counter available drug used to treat depression, obesity, insomnia and chronic headaches (Das et al., 2004)), as an analogous control compound derived from tryptophan, we screened for deamination of these compounds in batch cultures of *C. sporogenes*. Cultures were incubated with 100 μM levodopa or 5-HTP in combination with PAAAs from the growth medium and were followed over a period of 48 hours. Analysis of the samples using High Pressure Liquid Chromatography (HPLC) coupled to an electrochemical detector (ED) revealed that levodopa is completely converted within 24 h to a new metabolite, which was identified by ^1^H/^13^C-NMR and LC-MS as 3-(3,4-dihydroxyphenyl)propionic acid, DHPPA (**Figure 1B, 1C, Supplementary Figure 1A, 1B, 1C**). Furthermore, the incubations showed that the PAAAs available from the growth medium did not prevent the deamination of levodopa and that, during the incubation for 48 h, DHPPA remained stable. Similarly, 5-HTP was converted into two new unknown peaks (**Supplementary Figure 2A, 2B**), albeit to a much lesser extent compared to levodopa. Only the first peak could be detected and assigned by LC-MS as 5-hydroxyindole-3-lactic acid (5-HILA) by its predicted exact mass (**Supplementary Figure 2C**). The other peak is potentially 5-hydroxyindole-3-propionic acid (5-HIPA), described below.

**Figure 1.**
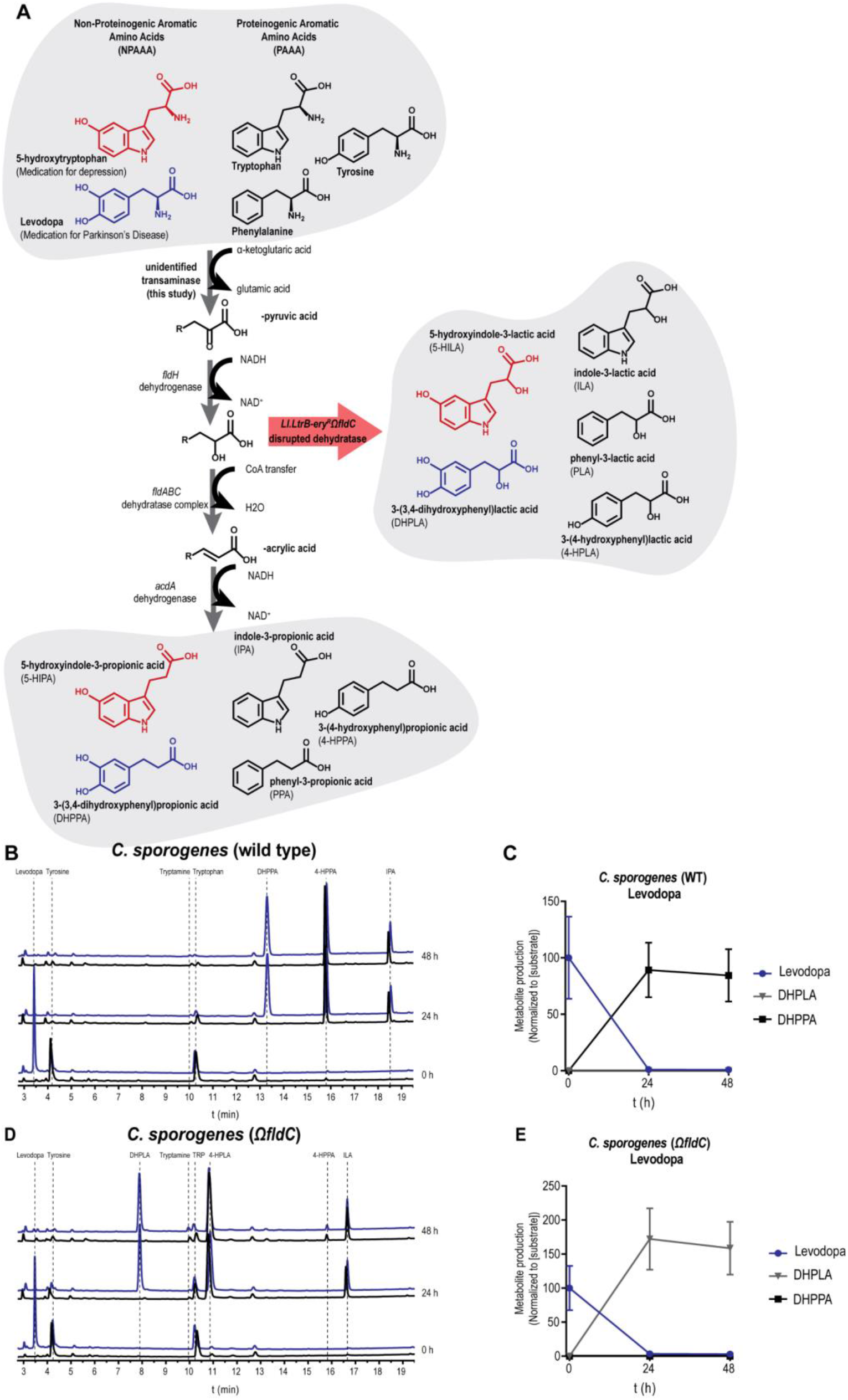
Levodopa is deaminated by *Clostridium sporogenes*. (**A**) Full reductive deamination pathway of *C. sporogenes* is depicted resulting in the full deamination (R-propionic acid) of (non)-proteinogenic aromatic amino acids (N)PAAA. The red arrow indicates a disrupted deamination pathway of *C. sporogenes*, where the dehydratase subunit *fldC* is mutagenized, resulting in a pool of partially deaminated metabolites (R-lactic acid) by *C. sporogenes*. (**B**) HPLC-ED curves from supernatant of a *C. sporogenes* batch culture conversion of levodopa (3-(3,4-dihydroxyphenyl)alanine) overtime. At the beginning of growth (timepoint 0 h) 100 μM of levodopa (blue) is added to the culture medium, the black line in the chromatogram depicts the control samples. In 24 h, levodopa is completely converted to DHPPA (3-(3,4-dihydroxyphenyl)propionic acid), the deaminated product of levodopa. Other aromatic amino acids from the medium, tryptophan and tyrosine (which are detectable with ED), are converted to the deaminated products IPA (indole-3-propionic acid) and 4-HPPA (3-(4-hydroxyphenyl)propionic acid). (**C**) Quantification (n=3) of levodopa conversion to DHPPA by *C. sporogenes* wild type (also see **Supplementary Table 1**). (**D**) Analysis of the supernatant of CS^Ω*fldC*^ shows that levodopa is not deaminated to DHPPA but to its intermediate product DHPLA (3-(3,4-dihydroxyphenyl)lactic acid) within 24 h. Tryptophan and tyrosine are converted to their intermediates ILA (indole-3-lactic acid) and 4-HPLA (3-(4-hydroxyphenyl)lactic acid), respectively. (**E**) Quantification (n=3) of levodopa conversion to DHPLA by *C. sporogenes* Ω*fldC* (also see **Supplementary Table 1**). All experiments were performed in 3 independent biological replicates and means with error bars representing the SEM are depicted.

To further investigate the involvement of the deamination pathway in levodopa and 5-HTP deamination, the enzyme responsible for the dehydratase reaction (encoded by the *fldC* gene (Dickert et al., 2000, 2002; Dodd et al., 2017)) was disrupted using the ClosTron mutagenesis system (**Supplementary Figure 2D**) (Heap et al., 2010). The resulting strain *C. sporogenes Ll.LtrB-ery^R^ΩfldC* (CS^Ω*fldC*^) was incubated with levodopa, and the PAAAs from the growth medium. Tryptophan and tyrosine were converted to their intermediates ILA (indole-3-lactic acid) and 4-HPLA (3-(4-hydroxyphenyl)lactic acid), respectively, as previously shown (Dodd et al., 2017). Analogous to tryptophan and tyrosine, levodopa was no longer deaminated to DHPPA but to its intermediate product 3-(3,4-dihydroxyphenyl)lactic acid (DHPLA) (**Figure 1D, 1E**). Only a slight production of 4-HPPA (from tyrosine) is observed after 48 h, presumably because of the substitution of FldABC by the similar HadABC proteins from the *had*-operon in *C. sporogenes* (Dickert et al., 2002; Dodd et al., 2017). HPLC-ED analysis of the 5-HILA production from 5-HTP by the *fldC* mutant was hampered by the production of coeluting 4-HPLA, the intermediate deamination product produced from tyrosine (described above). However, the analysis revealed that the second unknown peak produced from 5-HTP was no longer produced by CS^Ω*fldC*^ (**Supplementary Figure 2E, 2F**), demonstrating that 5-HTP conversion is affected and suggesting that the unknown product is 5-hydroxyindole-3-propionic acid (5-HIPA). Overall, the results show that the deamination pathway from *C. sporogenes* is not only involved in the deamination of PAAAs but also is in the deamination of the NPAAAs, levodopa and 5-HTP.

### Identification of the aromatic aminotransferase responsible for initiation of the deamination pathway

The aromatic aminotransferase responsible for the transamination of levodopa and the other (N)PAAAs, is crucial for the initiation of the reductive deamination pathway and for the full deamination of the substrates by the dehydrogenases (FldH and AcdA) and dehydratase (FldABC) (**Figure 1A**). However, the gene encoding this transaminase remains unidentified. To further investigate this critical step in the pathway, all nine class I/II aminotransferases encoded by *C. sporogenes* were cloned, purified and screened for their activity on levodopa and the other (N)PAAAs. Screening revealed a single aminotransferase (EDU38870 encoded by CLOSPO_01732) to be involved in their transamination (**Figure 2A**). To verify whether other aminotransferases could substitute for the identified aminotransferase *in vivo*, CLOSPO_01732 was disrupted (resulting in CS^Ω*CLOSPO_01732*^ (**Supplementary Figure 3A**)) and a targeted metabolomic analysis of all the (N)PAAA metabolites was performed using HPLC-ED (except metabolites from phenylalanine, which were quantified using HPLC-UV). The disruption of *fldC* or *CLOSPO_01732* resulted in only a minor reduction of the exponential growth rate in rich broth (Doubling time is 55.1±1.2 min and 64.1±1.1 min respectively compared to wild type 44.3±1.2 min) all reaching stationary phase within 12 hr (**Supplementary Figure 3B**). Comparing the metabolic profiles from wild type *C. sporogenes* (CS^WT^), CS^*ΩfldC*^ and CS^*ΩCLOSPO_01732*^ demonstrated that none of the other tested aminotransferases could take over this transaminase reaction effectively, except for the substrate phenylalanine (**Figure 2B and Supplementary Table 1**). Disrupting CLOSPO_01732 significantly reduced the production of phenyl-3-propionic acid (PPA), 3-(4-hydroxyphenyl)propionic acid (4-HPPA), indole-3-propionic acid (IPA), and 3-(3,4-dihydroxyphenyl)propionic acid (DHPPA) by 16.4%, 79.0%, 97.2%, 97.7%, respectively compared to CS^WT^ within 24-48 h (**Supplementary Table 1**). Presumably, the transamination of phenylalanine is substituted by EDU37030 as this aminotransferase also showed phenylalanine-converting activity *in vitro* (**Figure 2A**). Interestingly, CS^*ΩCLOSPO_01732*^ produces significantly higher amounts of tryptamine (4 to 6-fold increase at 24 and 48 h, respectively) compared to CS^WT^, reflecting a reduced competition for the same substrate by different enzymes (**Figure 2B**, **Supplementary Table 1**). Analogous to tryptamine, CS^*ΩCLOSPO_01732*^ produced significantly more serotonin compared to CS^WT^ at 48 h when incubated with 5-HTP (**Figure 2B**, **Supplementary Figure 3C, Supplementary Table 1**), though to a much lesser extent (~1% of substrate added) compared to tryptamine. Collectively, the data show that the aromatic aminotransferase (EDU38870), is involved in the initiation of the aromatic amino acid deamination pathway and is crucial for the production of DHPPA, 5-HILA, 5-HIPA, and as well as the previously described metabolites to be circulating in the blood, IPA, and 4-HPPA (Dodd et al, 2017).

**Figure 2.**
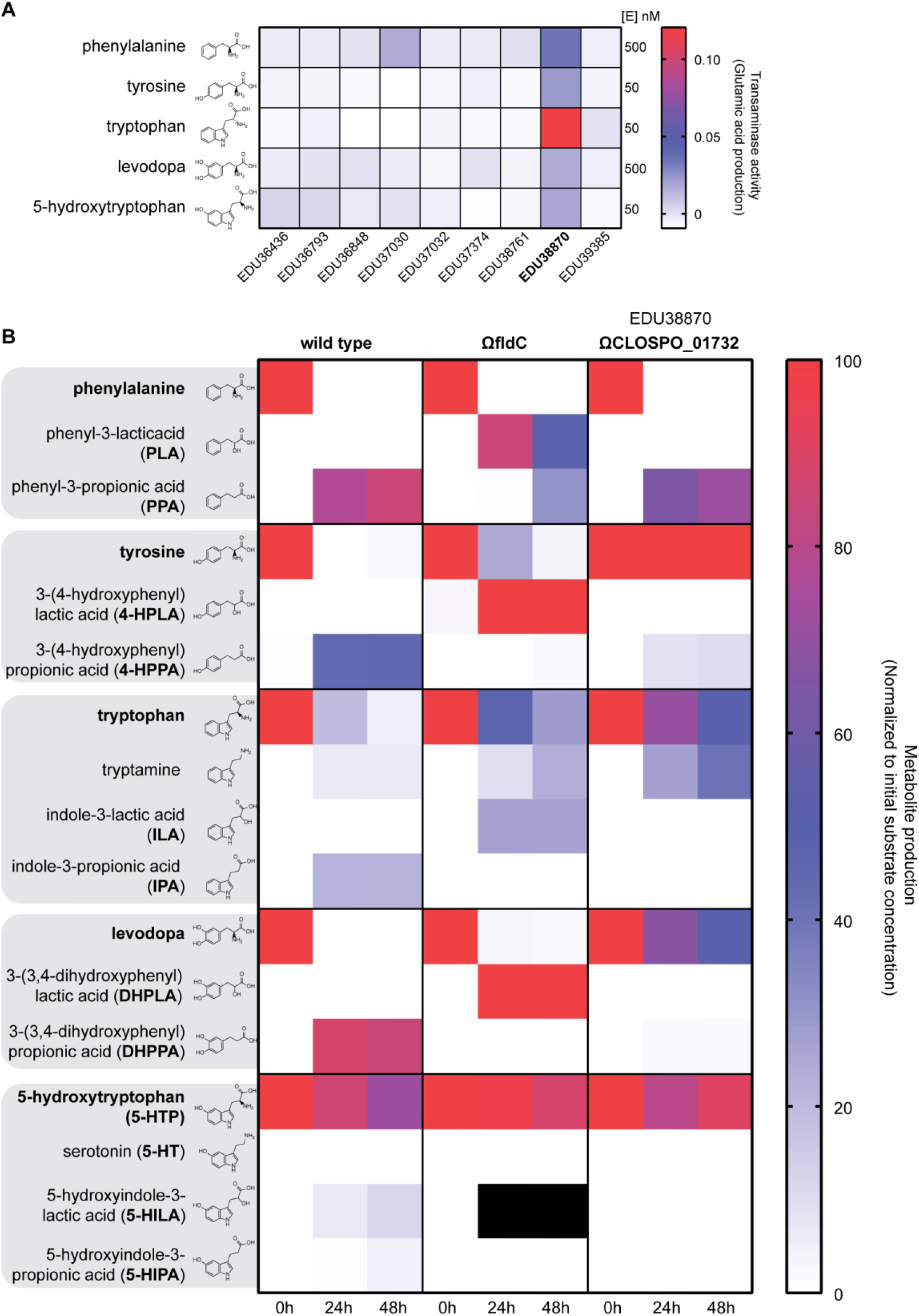
Identification of the aromatic amino transferase initiating the deamination pathway. In order to identify which aminotransferase is responsible for the initial transaminase reaction all class I/II aminotransferase were cloned and purified to test the activity against (N)PAAAs. (**A**) Transaminase activity (production of glutamate) for all substrates is depicted. EDU38870 (CLOSPO_01732) was involved in all transaminase reactions. EDU37030 showed similar activity as EDU38870, for phenylalanine. Experiment was performed in technical duplicates to screen for candidate genes for mutagenesis in *C. sporogenes*. (**B**) Targeted metabolic quantification of deamination products from CS^WT^, CS^ΩfldC^, CS^ΩCLOSPO_01732^ reveals that EDU38870 is involved in the transamination of all for all tested (N)PAAAs. All quantified deamination products are normalized to their initial substrate concentration and the data represents 3 independent biological replicates. Corresponding values are reported and metabolite concentration differences between WT and *ΩfldC* or ΩCLOSPO_01732 were tested for significance using Student’s t-Test, in **Supplementary Table 1**. Black squares indicate that quantification was not possible because of a coeluting HPLC-ED peak. As no commercial standards are available for 5-HILA and 5-HIPA, the peaks were quantified assuming a similar ED-detector response as for 5-HTP.

### 3-(3,4-dihydroxyphenyl)propionic acid elicits an inhibitory effect on ileal muscle contractions *ex vivo*

Because levodopa is the main treatment of PD patients, and is efficiently deaminated to DHPPA within 24 h by the *C. sporogenes* deamination pathway compared to 5-HTP, we further focused on levodopa and its deamination products. DHPPA is a phenolic acid (a molecule in the class polyphenols) and recent findings demonstrated an association between bacterial-derived polyphenol metabolites and gut-transit times in humans (Roager et al., 2016). Levodopa is mainly absorbed in the proximal small intestine, but significant amounts can reach the distal part of the intestinal tract (Morgan, 1971), and these levels increase with age (Iwamoto et al., 1987). As levodopa is taken orally, the first intestinal site where anaerobic bacteria such as *C. sporogenes* (*Clostridium* Cluster I) can encounter relevant levels of levodopa is the ileum. Studies on asymptomatic ileostomy subjects established that the core ileal microbiota consists of (facultative) anaerobes including species from *Clostridium* Cluster I (Booijink et al., 2010; Zoetendal et al., 2012). Moreover, the transit time in the SI has been shown to be significantly longer in PD patients compared to healthy controls (with a median increase of 1.75 hours in PD patients) (Dutkiewicz et al., 2015; Knudsen et al., 2017). To this end, we tested whether DHPPA (100 μM) could affect the muscle contractility in the ileum. Ileal rings of wild type C57BL/6J mice were suspended in an *ex vivo* organ bath system to test the effect of DHPPA on muscle contractions. Our initial results indicated that DHPPA displayed an inhibiting effect on natural ileal contractility (**Supplementary Figure 4A**).

Because acetylcholine is the neurotransmitter constantly produced from the excitatory muscle motor neurons to induce gut smooth muscle contractility (Costa et al., 2000), we tested whether DHPPA could have an inhibiting effect on acetylcholine-induced contractility in the ileum. The differences in amplitude of the contractions were quantified by measuring the decrease of the observed frequencies after a Fourier transform of 5 min intervals (**Figure 3A**). Ileal tissue preparations were tested by initiating an acetylcholinergic twitch by adding 50 μM of acetylcholine (a concentration saturating the muscarinic receptors (Kd=1.7±0.18 μM (Ringdahl, 1986)). After five minutes, 100 μM DHPPA (a concentration resembling the higher levels detected in fecal samples of PD patients, see below) was added and contractions were followed further over a period of 15 minutes. One-minute traces of the contractility representing one of the experiments are shown before and after addition of acetylcholine, DHPPA or vehicle (**Figure 3B**). A significant decrease in the amplitude (binned in 5 min intervals) of the acetylcholinergic twitch by DHPPA was observed at the 10-15 min (maximal reduction 69%) and 15-20 min interval (maximal reduction 73%) (**Figure 3C**). In order to determine the potency of DHPPA, a dose response curve with DHPPA was performed and showed a half maximal inhibitory concentration (IC_50_) of 20.3±10.6 μM (**Figure 3D**). In contrast to DHPPA, incubations with levodopa did not show any significant effect on the acetylcholinergic twitch (**Supplementary Figure 4B).** Collectively the data shows that DHPPA can inhibit the acetylcholine-induced muscle contractility of mouse ileum *ex vivo*.

**Figure 3.**
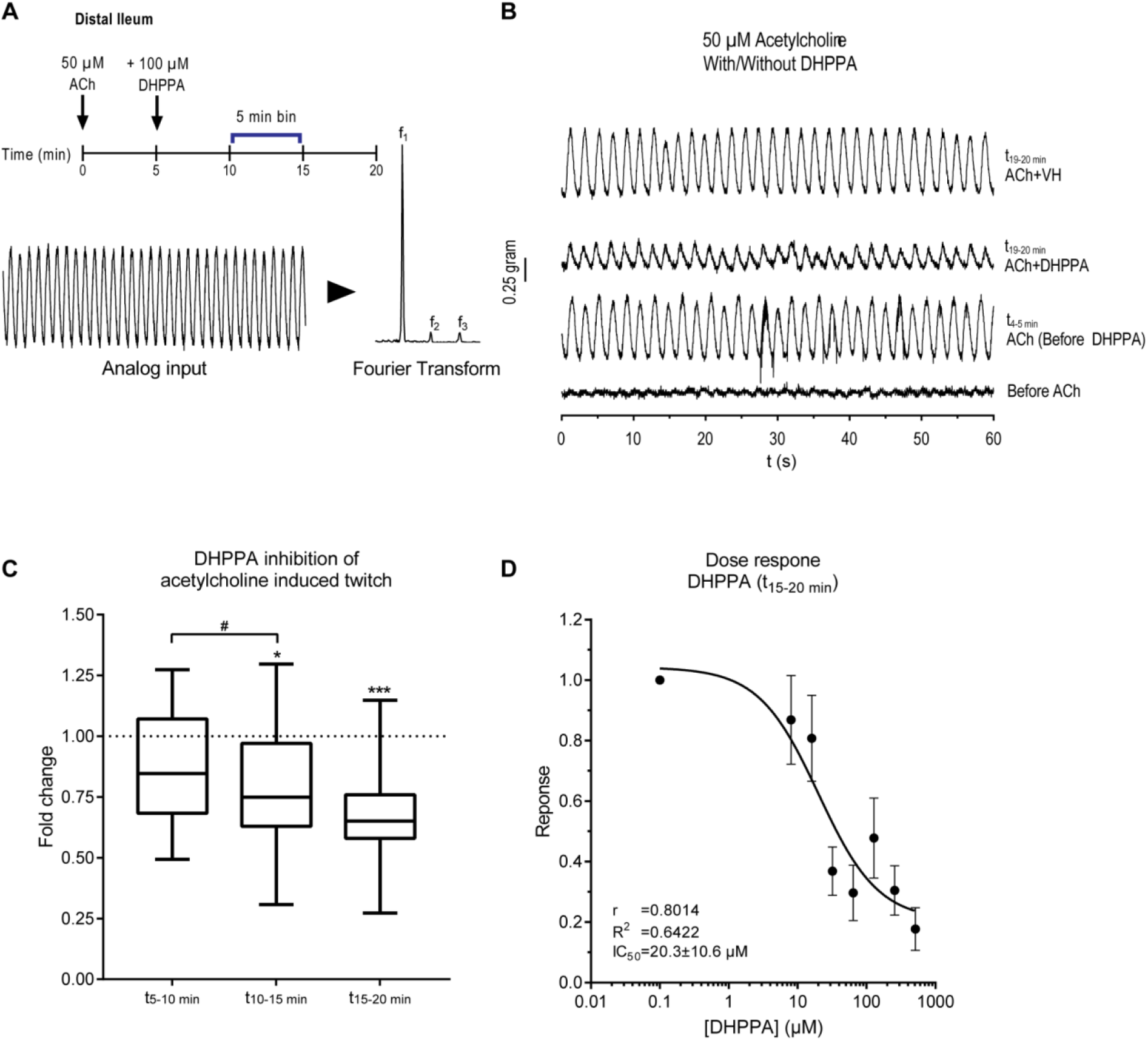
DHPPA inhibits the acetylcholine-induced twitch from mouse ileum. (**A**) Experimental setup, where 5 min after adding 50 μM acetylcholine, 100 μM DHPPA is added. The panel below indicates how the amplitude of the frequencies of the observed oscillations (from 5 minute bins) are extracted by a Fourier transform of the analog input. (**B**) A representative 1 min recording trace before and after the addition of acetylcholine and DHPPA or vehicle (VH) is shown. ACh, acetylcholine; VH, vehicle (0.05% ethanol). (**C**) Inhibition of DHPPA on acetylcholine-induced twitch binned in intervals of 5 minutes shows a decrease in contractility over time (n=6 biological replicates and experiments were repeated 1-4 times per tissue). Significance was tested using repeated measures (RM) 1-way-ANOVA followed by a Tukey’s test (*=p<0.0021 ***=p<0.0002 #=p<0.0021). Box represents the median with interquartile range and whiskers represent the maxima and minima. (**D**) Dose response curve of DHPPA on the acetylcholine-induced twitch at the t_15-20_ minute bin (n=4 biological replicates) with a half maximal inhibitory concentration (IC_50_) of 20.3±10.6 μM.

### Active levodopa deamination pathway in fecal suspensions of patients with Parkinson disease

We hypothesized that if *C. sporogenes* or other bacteria with the deamination pathway (*C. botulinum, Peptostreptococcus anaerobius* or *Clostridium cadaveris* (Dodd et al., 2017)) are present in the GI-tract of PD patients on levodopa/carbidopa treatment, those patients might have considerable amounts of DHPPA in their distal GI-tract. Because DHPPA can be a product of gut bacterial metabolism of polyphenolic rich foods in the colon such as coffee and fruit (Jenner et al., 2005), fecal samples from 10 PD patients were compared to 10 age-matched HC. Samples were collected in a previous study and there were no significant difference in in macronutrients, dietary fiber, or total calorie intake between groups (Keshavarzian et al., 2015). Using a catechol extraction targeted for the quantification of DHPPA we found that the DHPPA concentrations were significantly higher in PD patient’s fecal samples compared to HC (Figure 4A). Identification of DHPPA was confirmed by LC-MS (Supplementary Table 2). The higher amounts (2.2 fold increase) of DHPPA observed in the fecal samples of PD patients are likely to result from levodopa metabolized by the anaerobic bacteria, deaminating levodopa through the FldBC dehydratase (Figure 1A). In order to investigate the presence and activity of the anaerobic deamination pathway in fecal samples, the dehydratation of the intermediate levodopa metabolite, DHPLA (Figure 1A) was tested. The levodopa intermediate DHPLA was used as substrate instead of levodopa to prevent an *in vitro* substrate bias for bacteria that can decarboxylate levodopa to dopamine (van Kessel et al., 2019; Maini Rekdal et al., 2019). Moreover, FldABC is the key protein complex responsible for the production of DHPPA. Screening for the identified transaminase or FldH dehydrogenase upstream of FldABC would not be relevant as many bacterial species harbor these type of enzymes (Supplementary Figure 5). Hence, fecal suspensions (10% w/v) from PD and HC were incubated anaerobically with DHPLA, and samples were collected at 0, 20, and 45 h and were analyzed by HPLC-ED. After 20 h, DHPPA was detected in fecal samples from PD patients, as well as in fecal samples of HC when supplied with the substrate levodopa (**Figure 4B, Supplementary Figure 6A**). Moreover, DHPPA was further converted to the downstream dehydroxylated metabolite of DHPPA, 3-(3-hydroxyphenyl)propionic acid (3-HPPA) over time (**Figure 4B, Supplementary Figure 6A**). Because DHPPA is further converted to 3-HPPA *in vitro* we quantified both the production of DHPPA and/or 3-HPPA in the fecal incubations as measure for the presence of an active deamination pathway. Metabolic profiles of PD or HC samples that produced DHPPA/3-HPPA overtime were quantified and merged (**Figure 4C, Supplementary Figure 6B**), showing that DHPPA is produced first, and is further metabolized to 3-HPPA. The production of DHPPA or 3-HPPA was observed in 50% and 20% of the PD patient’s and HC fecal-suspensions, respectively after 20 h and in 70% and 50% PD patient’s and HC fecal-suspensions, respectively after 45 h (Supplementary Figure 6C). The production of 3-HPPA *in vitro* is likely to be performed by *Eggerthella lenta*, which has been shown to perform *p*-dehydroxylations (Jin and Hattori, 2012). Indeed *in vitro* culturing of *E. lenta* showed *p*-dehydroxylation of DHPPA (Supplementary Figure 7A). Because DHPPA is further converted to 3-HPPA *in vitro,* we examined whether 3-HPPA could elicit a similar effect on the acetylcholine induced contractions in ileum. Unlike, DHPPA, 3-HPPA did not elicit a significant effect on the acetylcholine induced twitch (Supplementary Figure 7B). Furthermore, to investigate the genomic abundance levels of bacteria capable of deaminating (N)PAAAs, we analyzed the 16s rDNA sequence data of the fecal samples of patients with Parkinson disease (Keshavarzian *et al.,* 2015) that were employed in this study (**Supplementary Results and Supplementary Figure 8**). A significant positive correlation (*r*= 0.62, R^2^= 0.38, *p*= 0.02) was found between bacteria with the deamination pathway and DHPPA/3HPPA production in fecal incubation samples at 20 h (**Supplementary Figure 8E**). Taken together, the results show that DHPPA can be produced by the microbiota via anaerobic deamination of levodopa. Moreover, our findings indicate that 3-HPPA originates from DHPPA via dehydroxylation potentially by *Eggerthella lenta* and that the aromatic deamination pathway, as measured by the production of DHPPA or 3-HPPA, is active and present in at least 70% of the PD samples.

**Figure 4.**
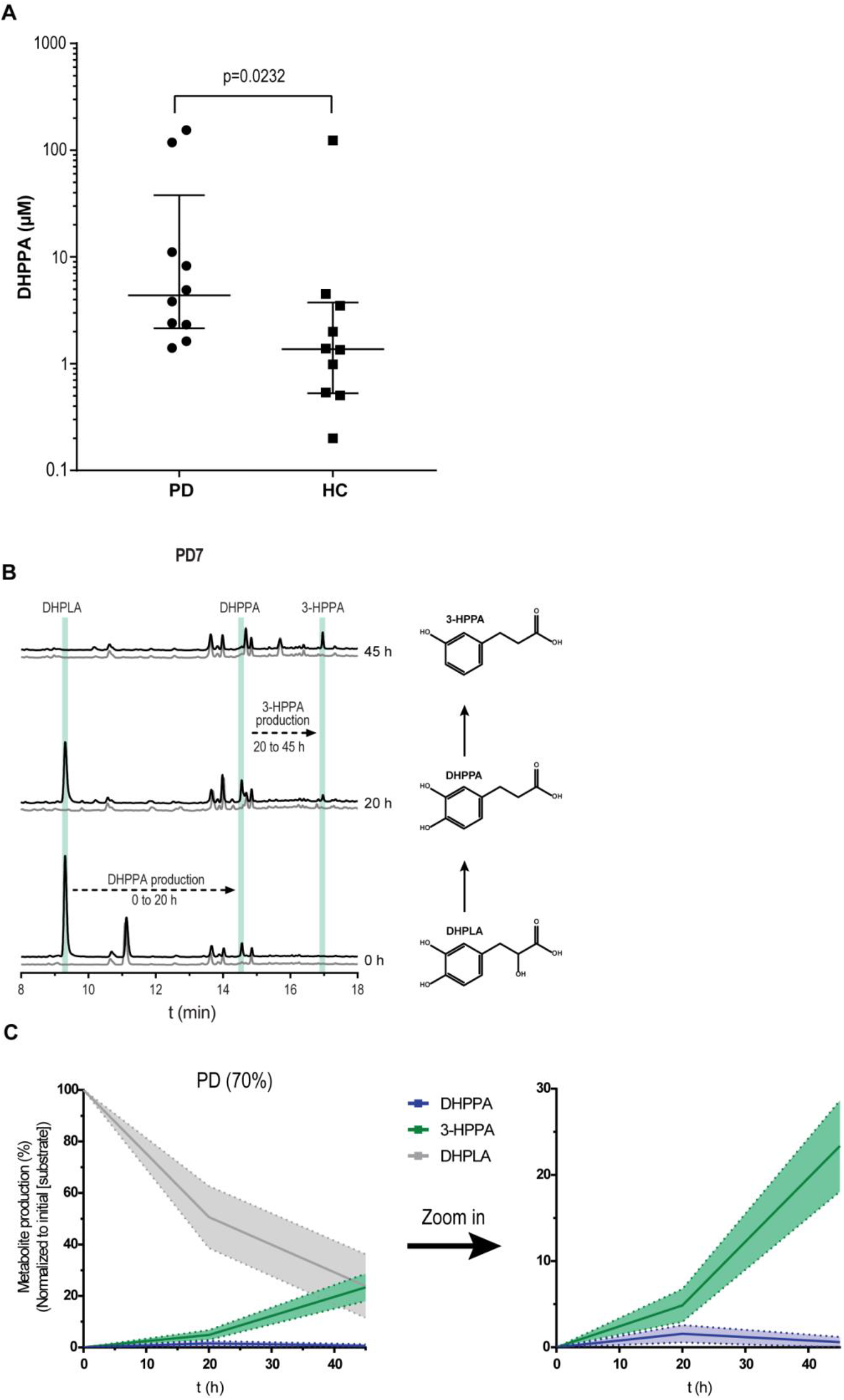
Higher DHPPA levels in PD patients and active levodopa deamination pathway in PD fecal suspensions. (**A**) DHPPA was extracted from fecal samples of PD patients (n=10) and age-matched healthy controls (n=10) using activated alumina beads and concentrations were quantified using a standard curve of DHPPA on the HPLC-ED with 3,4-dihydroxybenzylamine as internal standard. DHPPA concentrations are depicted on the logarithmic y-axis and individual levels are indicated and compared between Parkinson’s disease patients (PD) and age-matched healthy controls (HC). The cross-header represents the median (PD, 4.36 μM; HC, 1.37 μM) and the interquartile range (PD, 2.15-37.90 μM; HC, 0.53-3.75 μM). Significance was tested using an unpaired nonparametric Mann-Whitney test (p=0.0232) (**B**) A representative HPLC-ED chromatogram of fecal-suspension from PD7 where DHPPA is produced from DHPLA (black) after 20 h and is further metabolized to 3-HPPA after 45 h of incubation. The control, without the addition of DHPLA is indicated in grey. The green bars indicate the retention time of the standards indicated. (**C**) Metabolite profiles of the PD fecal suspensions that produced DHPPA/3-HPPA within 20-45h (70%) are merged as replicates. Lines represent the mean and the shadings the SEM, a zoom in graph of DHPPA and 3-HPPA is depicted on the right.

## Discussion

Identifying bacterial pathways and elucidating their potential impact on bacterial drug metabolism is crucial in order, not only, to maximize medication efficacy, but also to recognize and eventually prevent potential side effects that might affect the host’s physiology on an individual basis (Haiser et al., 2013; van Kessel et al., 2019; Niehues and Hensel, 2009; Zimmermann et al., 2019). Here we disclosed the reductive anaerobic deamination pathway in *C. sporogenes* by identifying its initiating enzyme, the aromatic aminotransferase, and expanded the pathway’s relevance by demonstrating its capacity to convert two clinically important NPAAAs, levodopa and 5-HTP. We showed that *C. sporogenes* is able to completely deaminate levodopa to DHPPA and to a much lesser extent 5-HTP (**Figures 1, 2 Supplementary Figure 1, 2, 3 and Supplementary Table 1**). Disrupting the bacterial transaminase encoding gene abolished the production of deaminated products, and increased the production of neuromodulators such as tryptamine and, to much lesser extent, serotonin (**Figure 2, Supplementary Figure 3C and Supplementary Table 1**). Tryptamine is a natural product produced by *C. sporogenes* that has been proposed to modulate gut transit time (Williams et al., 2014). The application of engineered gut bacteria as a therapeutic strategy to modulate GI-motility or host physiology has been also proposed recently in two proof-of-concept studies; by heterologous expression of *Ruminococcus gnavus* tryptophan decarboxylase in *Bacteroides thetaiotamicron* (Bhattarai et al., 2018) and by modification of the metabolic output of bioactive compounds in an engineered *fldC* deficient *C. sporogenes* strain (Dodd et al., 2017). However, translation of these studies into applications is hindered by restrictions on the application of genetically engineered microorganisms (GMOs) *per se*, and the complexity of introducing these GMOs into an existing gut microbiota ecosystem. Selective therapeutic blockage of the aminotransferase identified in this study may provide an attractive alternative solution to modify gut microbiota metabolism.

PD patients encounter increased gut transit time; thus, an additional inhibition of acetylcholine-induced contraction could result in further slowing down of gut transit rates. The inhibitory effect of DHPPA on the acetylcholine-induced ileal muscle contractions (**Figure 3**), higher DHPPA levels in fecal samples of PD patients compared to HC (**Figure 4**), and an active deamination pathway of levodopa during fecal-incubations of PD patients (**Figure 4**) demonstrate active deamination of levodopa in the distal GI-tract of PD patients and suggest potential side effects of this bacterial by-product of the unabsorbed residue of the medication. DHPPA shares similarity with dopamine structure except of the terminal amine group, which is substituted by a carboxyl group in DHPPA. Dopamine, and dopamine agonists, have been shown to inhibit methacholine (analog of acetylcholine) induced contraction, which is not mediated via dopamine receptors, in guinea pig jejunum in similar concentration ranges to DHPPA (EC_50_ relaxation by dopamine ~290 μM) (Lucchelli et al., 1990), indicating that DHPPA might act on a similar mechanism. Although further research is needed to unravel the underlying mechanism, these results indicate that DHPPA inhibits the acetylcholine-induced muscle contractions in the ileum with implications on intestinal motility, often observed in PD patients. Overall, the present study highlights the urgency to unravel potential effects of gut bacterial processing of (unabsorbed residues of) medication, such as levodopa.

## Material and Methods

### Growth and incubation of *Clostridium sporogenes* and *Eggerthella lenta*

*Clostridium sporogenes* ATCC15579 was grown in enriched beef broth (EBB) with 2 g/L glucose (van Kessel et al., 2019) and 0.1% Tween 80 (EBB/T) anaerobically (10% H_2_, 10% CO_2_, 80% N_2_) in a Don Whitley Scientific DG250 Workstation (LA Biosystems, Waalwijk, The Netherlands) at 37 °C. *Eggerthella lenta* DSM2243 was grown in modified DSMZ medium 78 (DSMZ 78: Beef extract, 10.0 g/L; Casitone, 30.0 g/L; Yeast extract, 5.0 g/L; K2HPO4, 5.0 g/L; Tween 80, 0.1%; Menadione (Vitamin K3), 1 μg/ml; Cysteine, 0.5 g/L; Hemin, 5 mg/L; L-Arginine 0.1-1.5%) anaerobically (1.5% H_2_, 5% CO_2_, balance with N_2_) in a Coy Laboratory Anaerobic Chamber (neoLab Migge GmbH, Heidelberg, Germany) at 37 °C in a tube shaker at 500 RPM. Upon use, bacteria were inoculated from −80 °C stored glycerol stocks in the appropriate media and grown for 18-24 h for *C. sporogenes* and 24-40 h for *E. lenta*. Overnight turbid cultures were then diluted 1/50 in an appropriate volume EBB/T or CMM for further experiments with 100 μM levodopa (D9628, Sigma), 5-hydroxytrytophan (H9772, Sigma), 50 μM 3-(3,4-dihydroxyphenyl)propionic acid (102601, Sigma) or H_2_O as control. All experiments were performed in triplicate (3 biological replicates).

### Protein production and purification

Transaminase-encoding genes from *C. sporogenes* (**Supplementary Table 3**) were amplified using Phusion High-fidelity DNA polymerase and primers listed in **Supplementary Table 3**. All amplified genes were cloned in pET15b, except for EDU37032 which was cloned in pET28b (**Supplementary Table 3**). Plasmids were maintained in *E. coli* DH5α and verified by Sanger sequencing before transformation to *E. coli* BL21 (DE3). Overnight cultures were diluted 1:50 in fresh LB medium with the appropriate antibiotic and grown to OD600 = 0.7–0.8 shaking at 37 °C. Protein translation was induced with 1 mM Isopropyl β-D-1-thiogalactopyranoside (IPTG, 11411446001, Roche Diagnostics) and cultures were incubated overnight at 18 °C. The cells were washed with 1/5th of the volume in 1× ice-cold PBS and stored at −80 °C or directly used for protein isolation. Cell pellets were thawed on ice and resuspended in 1/50th of buffer A (300 mM NaCl; 10 mM imidazole; 50 mM KPO4, pH 8.0) containing 0.2 mg/mL lysozyme (105281, Merck) and 2 μg/mL DNAse (11284932001, Roche Diagnostics), and incubated for at least 10 min on ice before sonication (10 cycles of 15 s with 30 s cooling at 8 microns amplitude) using Soniprep-150 plus (Beun de Ronde, Abcoude, The Netherlands). Cell debris was removed by centrifugation at 20,000 × g for 20 min at 4 °C. The 6 × his-tagged proteins were purified using a nickel-nitrilotriacetic acid (Ni-NTA) agarose matrix (30250, Qiagen). Cell-free extracts were loaded on 0.5 ml Ni-NTA matrixes and incubated on a roller shaker for 2 h at 4 °C. The Ni-NTA matrix was washed three times with 1.5 mL buffer B (300 mM NaCl; 20 mM imidazole; 50 mM KPO4, pH 8.0) before elution with buffer C (300 mM NaCl; 250 mM imidazole; 50 mM KPO4, pH 8.0). Imidazole was removed from purified protein fractions using Amicon Ultra centrifugal filters (UFC505024, Merck) and washed three times and reconstituted in buffer D (50 mM Tris-HCl; 300 mM NaCl; pH 7.5). Protein concentrations were measured spectrophotometrically (Nanodrop 2000, Isogen, De Meern, The Netherlands) using the predicted extinction coefficient and molecular weight from ExPASy ProtParam tool (www.web.expasy.org/protparam/).

### Transaminase activity test

Purified transaminases were incubated with 1 mM substrate, 2 mM α-ketoglutaric acid, and 0.1 mM PLP (pyridoxal-5-phosphate, P9255, Sigma, The Netherlands) in buffer D with an enzyme concentration of 50 nM for tyrosine, tryptophan, or 5-HTP as substrate and an enzyme concentration of 500 nM for phenylalanine and levodopa as substrate. Enzyme reactions were incubated for 0.5 h at 37 °C, the reactions were stopped with 0.7% (v/v) perchloric acid (1:1). Transaminase activity was tested using an L-glutamic acid detection kit (K-GLUT, Megazyme Inc., Wicklow, Ireland), according to the manufacture’s microplate assay procedure with some modifications. The supplied buffer was substituted for buffer D (described above, to prevent oxidation of the substrates/products). A reaction mix was prepared mixing 50 μL buffer D; 10 μL quenched sample reaction mixture, 20 μL NAD^+^/iodonitrotetrazolium chloride solution, 5 μL diaphorase solution, 5 μL glutamate dehydrogenase (GIDH) solution and reconstituted to a final volume of 290 μL with H2O. Absorbance at 492 nm was measured after 10 minutes of incubation using a microplate reader (Synergy HTX spectrophotometer, BioTek, BioSPX, The Netherlands) and background was subtracted from initial read before addition of GIDH solution.

### Targeted mutagenesis

Gene disruptions in *Clostridum sporogenes* were performed using the ClosTron system (Heap et al., 2007, 2009). This system facilitates targeted mutagenesis using the group-II *Ll.LtrB* intron of *Lactococcus lactis*. Introns targeting *fldC* (CLOSPO_311) or CLOSPO_1732 (encoding for the transaminase) were designed using the ClosTron intron design tool (http://www.clostron.com) and were ordered in pMTL007C-E2 from ATUM (Newark, California, United States) resulting in pMTL007C-E2_Cs-fldC-561a and pMTL007C-E2_Cs-CLOSPO_1732-493s respectively. Plasmids were transferred to *C. sporogenes* by conjugation as described before (Heap et al., 2009) using *E. coli* CA434 (*E. coli* HB101 (Bio-Rad Laboratories, The Netherlands) harboring the broad host IncPß^+^ conjugational plasmid pRK24 (Williams et al., 1990) as donor strain. *E. coli* CA434 harboring pMTL007C-E2_Cs-fldC-561a or pMTL007C-E2_Cs-CLOSPO_1732-493s was grown in Luria Broth (LB) with 10 μg/mL tetracycline and 25 μg/mL chloramphenicol (to select for pRK24 and PMTL007C-E2 respectively). Cell suspensions of 1 mL of overnight culture were washed once with PBS and the cell pellet was resuspended in 200 μL of *C. sporogenes* overnight cell suspension. The bacterial-mixture was spotted (in drops of 10 μL) on trypticase soy agar (TSA) plates and incubated for 24 h anaerobically at 37 °C. Sequentially, 1 mL of PBS was added to the spotted plates and the donor-recipient mix was scraped of the plate, sequentially the scraped-off suspension was distributed over TSA plates containing 50 μg/mL neomycin (to prevent growth of *E. coli*) and 15 μg/mL chloramphenicol to select for *C. sporogenes* conjugants. Chloramphenicol resistant colonies of *C. sporogenes* were re-streaked on TSA plates containing 50 μg/mL neomycin and 2.5 μg/mL erythromycin (to select for intron insertion) for several times. To makes sure the plasmids was integrated, colonies were checked and selected for their sensitivity towards chloramphenicol and the genomic DNA was verified using PCR (**Supplementary Figure 1F and 2A**)

### Fecal samples from patients with Parkinson’s disease and age-matched healthy controls

Fecal samples from patients diagnosed with PD (n = 10) and age-matched healthy controls (n = 10) were acquired from the Movement Disorder Center at Rush University Medical Center, Chicago, Illinois, USA published previously (Keshavarzian et al., 2015). All study subjects consented to the use of their samples for research. PD was diagnosed according to the UK Brain Bank Criteria as previously described (Keshavarzian et al., 2015). Study subjects were provided with the supplies and instructions for home feces collection using the BD Gaspak EZ Anaerobe Gas Generating Pouch System with Indicator (Ref 260683; Becton, Dickinson and Company, Sparks, MD) in order to minimize the exposure of the feces to high oxygen ambient atmosphere, which may alter the microbiota. Subjects were asked to have a bowel movement within 24 h of their study visit. Subjects kept the sealed anaerobic fecal bag in a cold environment, before bringing the anaerobic fecal bag to the hospital. Fecal samples were then immediately stored at −80°C until analysis.

### Fecal metabolite incubations from PD patients and HC subjects

Stool samples were suspended 1:1 (w/v) in EBB/T and incubated anaerobically (10% H_2_, 10% CO_2_, 80% N_2_) in a Don Whitley Scientific DG250 Workstation (LA Biosystems, Waalwijk, The Netherlands) at 37 °C with 100 μM Sodium 3-(3,4-dihydroxyphenyl)-DL-lactate (39363,Sigma). Samples were taken at 0, 20, and 45 h and analyzed on HPLC-ED as described below.

### HPLC-ED/UV analysis and sample preparation

For bacterial cell suspensions, 1 mL of methanol was added to 0.25 mL of cell suspension and stored at −20 °C until further use. For fecal metabolite incubations, 300 μl of methanol was added to 75 μL of fecal suspension and stored at −20°C until further use. Metabolites from stool samples were extracted by suspending the stool 1:1 (w/v) in water, followed by homogenization by vigorously vortexing while keeping samples as cold as possible. Homogenized suspensions were centrifuged at 3500 × *g* for 20 min at 4°C and sequentially 1.6 mL of methanol was added to 0.4 mL of supernatant. From bacterial, fecal incubation or stool samples cells and protein precipitates were removed by centrifugation at 20000 × *g* for 10 min at 4°C. Supernatant was transferred to a new tube and the methanol fraction was evaporated in a Savant speed-vacuum dryer (SPD131, Fisher Scientific, Landsmeer, The Netherlands) at 60°C for 1.5-2 h. The aqueous fraction was reconstituted with 0.7% HClO_4_ to the appropriate volume. Samples were filtered and injected into the HPLC-ED system (Alliance Separations Module 2695, Waters Chromatography B.V, Etten-Leur, The Netherlands; Dionex ED40 electrochemical detector, Dionex, Sunnyvale, USA, with a glassy carbon working electrode (DC amperometry at 0.8 or 1.0 V, with Ag/AgCl as reference electrode)). Samples were analyzed on a C18 column (Kinetex 5μM, C18 100 Å, 250 × 4.6 mm, Phenomenex, Utrecht, The Netherlands) using a gradient of water/methanol with 0.1% formic acid (0-10 min, 95-80% H_2_O; 10-20 min, 80-5% H_2_O; 20-23 min 5% H_2_O; 23-31 min 95% H_2_O). Fecal suspension metabolites were injected twice and analyzed at DC amperometry at 0.8 V (for DHPPA) and at 1.0 V (for 3-HPPA). Lowering the voltage makes the detection more selective for more readily oxidizable compounds (Nagatsu and Kojima, 1988) such as DHPPA, but making 3-HPPA invisible for detection. For the detection of the *C. sporogenes* metabolites and for peak isolation another HPLC-ED system was used (Jasco AS2059 plus autosampler, Jasco Benelux, Utrecht, The Netherlands; Knauer K-1001 pump, Separations, H. I. Ambacht, The Netherlands) with the same detector (ED40) and the same gradient as described above. Phenylalanine metabolites were detected by injecting the same samples in an HPLC-UV system (Alliance Separations Module 2695, Waters Chromatography B.V, Etten-Leur, The Netherlands; TSP UV6000LP UV-detector (wavelength: 260 nM) Thermo Scientific, The Netherlands). Samples for peak isolation were separated on a Vydac Semi-preparative C18 column (218TP510, 5 μm, 300 Å, 10 mm × 250 mm, VWR International B.V, Amsterdam, The Netherlands) at 3 ml/min using the same gradient as above. Data recording and analysis was performed using Chromeleon software (version 6.8 SR13). Significance was tested using a Two-sample equal variance (homoscedastic) Student’s t-Test (Microsoft Excel 2019 version 1808).

### Catechol extraction from stool for DHPPA quantification

Catechols were extracted from PD patients and HC stool samples using activated alumina powder (199966, Sigma) as previously described (van Kessel et al., 2019) with a few modifications. A volume of 200 μl 5O% stool suspension (described above) was used with 1 mM DHBA (3,4-dihydroxybenzylamine hydrobromide, 858781, Sigma) as an internal standard. Samples were adjusted to pH 8.6 with 800 μL TE buffer (2.5% EDTA; 1.5 M Tris/HCl pH 8.6) and 5–10 mg of alumina was added. Suspensions were mixed on a roller shaker at room temperature for 20 min and were sequentially centrifuged for 30 s at 20,000× *g* and washed three times with 1 mL of H_2_O by aspiration. Catechols were eluted using 0.7% HClO_4_ and filtered before injection into the HPLC-ED-system as described above (DC amperometry at 0.8 V). A standard curve was injected to quantify the concentrations of DHPPA in 50% (w/v) stool samples. Significance was tested using an unpaired nonparametric Mann-Whitney test (GraphPad Prism version 7).

### Organ-bath experiments

Distal ileal samples were harvested from wild type adult (18-20 weeks) male C57BL/6J mice that were sacrificed for another purpose. Harvested tissue was immediately removed, placed and washed in 0.85% NaCl. Approximately 3 mm rings were cut and were placed in an organ bath (Tissue Bath Station with SSL63L force transducer, Biopac Systems Inc. Varna, Bulgaria) filled with Krebs–Henseleit solution (NaCl, 7.02 g/L; KCl, 0.44 g/L; CaCl_2_.2H_2_O, 0.37 g/L; MgCl_2_.6H_2_O, 0.25 g/L; NaH_2_PO4.H_2_O 0.17 g/L; Glucose, 2.06 g/L; NaHCO_3_, 2.12 g/L) gassed with Carbogen gas mixture (5 % CO_2_, balanced with O_2_) at 37 °C. Ileal rings were equilibrated for at least 45-60 min with replacement of Krebs–Henseleit solution approximately every 15 min. Sequentially, 50 μM of acetylcholine (ACh) (Sigma, A2661) was added to induce a stable repetitive muscle twitch response, and after ~5 min 100 μM of DHPPA (102601, Sigma) (n=6 biological replicates, 1-4 technical replicates), 3-HPPA (91779, Sigma) (n=4 biological replicates, 2 technical replicates), or levodopa (D9628, Sigma) (n=3 biological replicates, 2 technical replicates) was added for ~15 min before the ileal rings were washed. This step was repeated 1-4 times per ileal preparation. As control, ACh was added for at least 20 min with or without 0.05% ethanol (solvent of DHPPA) after 5 min to check for spontaneous decrease. For the dose response curve (n= 4 biological replicates), every 15 minutes the cumulative dose of DHPPA was increased by 2-fold ranging from 8 to 512 μM. Data was recorded and analyzed in BioPac Student Lab 4.1 (Build: Feb 12, 2015). Frequencies were extracted performing a fast Fourier transform (FFT) on bins of 5 minute intervals. The maximum amplitude of all the observed frequencies were extracted and the average decrease of all frequencies overtime were calculated. Significance was tested using repeated measures (RM) 1-way-ANOVA followed by a Tukey’s test (GraphPad Prism version 7).

### Availability of data and materials

All data generated or analyzed during this study are included in this published article and its supplementary information files.

## Acknowledgements

We thank Mr. Walid Maho, Interfaculty Mass Spectrometry Center, University of Groningen, The Netherlands for running and analyzing the samples on the LC-MS and Prof. dr. Michiel Kleerebezem, Host-microbe interactomics group, Wageningen University, The Netherlands, for critical reading of our manuscript.

## Authors Contributions

S.P.K. and S.E.A conceived and designed the study. S.P.K., H.R.J., S.L.W., S.S.L., S.A.N., H.P., A.K. performed the experiments and S.P.K., S.S.L., H.P., and S.E.A. analyzed the data. S.P.K. and S.E.A. wrote the original manuscript that was reviewed by S.S.L., S.A.N., H.P., A.K. Funding was acquired by S.E.A.

