## Supplementary Information for "Gut bacterial deamination of residual levodopa medication for Parkinson’s disease"

### Supplementary Results

#### Analysis of deaminating bacteria and *E. lenta* in PD and HC fecal and mucosal samples

Measuring activity in fecal incubations shows that live bacteria express and produce enzymes that are capable of metabolic conversions, metagenomics on the other hand will only provide information whether a bacterium is present (death or alive) but no information on the activity of a certain metabolic pathway. Although its drawbacks it is of value to investigate the genomic abundance levels of bacteria capable of deaminating (N)PAAAs. In order to determine the relative abundance of deaminating bacteria, the 16s rDNA metagenomic sequence data from stool and sigmoid colon mucosa samples of PD patients and healthy controls from Keshavarzian *et al*., 2015 (bioproject PRJNA268515) were analyzed using Kraken2, a *k*-mer taxonomic classification system followed by Bracken (Bayesian Reestimation of Abundance with KrakEN) that computes the abundance of species. We extracted the bacteria that are known to be capable of deaminating (N)PAAAs (Dickert et al., 2000, 2002; Dodd et al., 2017) and *E. lenta* and compared their relative abundance between PD and HC samples.

In all fecal samples (prevalence = 1.0) *Clostridium botulinum* was detected. *E. lenta* was detected in 91% and 87% (prevalence = 0.91 and 0.87) of PD and HC samples, respectively. *C. sporogenes* was found in 2.9% of the PD samples only, although many *C. sporogenes* reads might be wrongly associated with the *C. botulinum* clade as, based on 16S rDNA, they are occurring in a single phylogenetic clade and some strains share high sequence similarity (≥ 99.8 %) of their 16S rRNA (Dickert et al., 2002; Kalia et al., 2011). Moreover, some *C. sporogenes* sequences show exact homology with *C. botulinum* based on 16S rDNA *in silico* restriction enzyme analysis  (Kalia et al., 2011). No reads were associated with *Clostridium cadaveris* or *Peptostrepotococcus anaerobius*. Furthermore, comparing the relative abundance of *C. botulinum* or *E. lenta* between PD and HC, no significant differences were observed between PD and HC fecal or mucosal samples (**Supplementary Figure 8A-D**), which is in agreement with the observed similar activity in PD and HC samples (**Supplementary Figure 6C**).

In order to investigate whether the DHPPA/3HPPA production in the fecal incubations are associated with higher levels of *C. botulinum* a correlation analysis was performed. The analysis showed a significant positive correlation ($r$= 0.62, $R^2$= 0.38, $p$= 0.02) between the relative abundance of *C.*

*botulinum* and DHPPA/3HPPA production in fecal incubation samples at 20 h (**Supplementary Figure 8E**). No significant correlation between the DHPPA levels extracted from the PD samples (**Figure 4A**) and *C. botulinum* was observed ($r$=-0.05, $R^2$=0.003, $p$=0.89), indicating that some DHPPA might have originated from other sources, which is in agreement with the fact that DHPPA is also observed in the HC samples (**Figure 4A**).

### Supplementary Figures

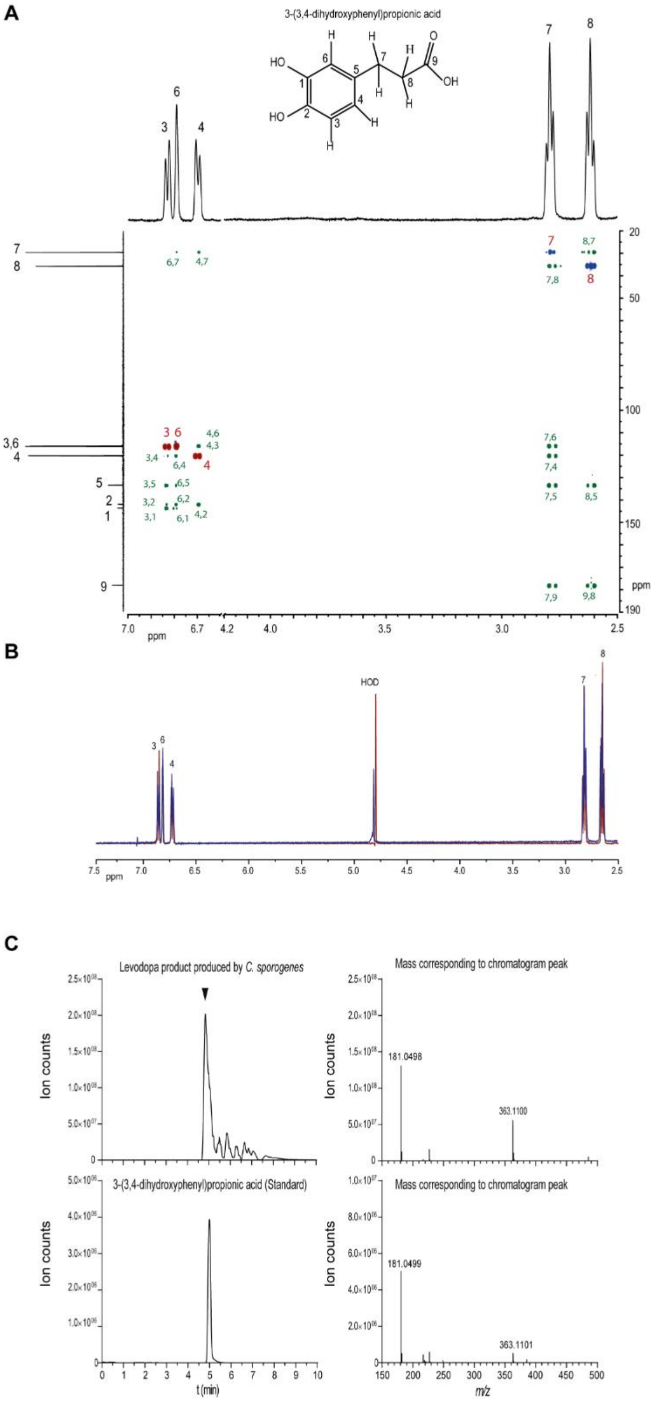

**Supplementary Figure 1. NMR and MS confirmation of levodopa product, 3-(3,4-dihydroxyphenyl)propionic acid. (A)** The isolated L-DOPA product was analysed by $^{1}$H and $^{13}$C NMR spectroscopy. The 1D $^{1}$H NMR spectrum showed 5 distinct peaks; δ 6.86 (*d J* 8.2 Hz; 1; H-3), δ 6.82 (*s*; 1; H-6), δ 6.72 (*d J* 8.6 Hz; 1; H-4 ), δ 2.82 (*t J* 7.2 Hz; 2; H-7), δ 2.64 (*t J* 7.2 Hz; 2; H-8). The 1D $^{13}$C NMR spectrum showed 9 peaks δ 178.47 (C-9), δ 143.80 (C-1), δ 142.08 (C-2), δ 133.69 (C-5), δ 120.46

(C-4), δ 116.26 (C-3), δ 116.07 (C-6), δ 35.80 (C-8) and δ 26.63 (C-7). The 2D $^{1}$H-$^{13}$C gHSCQ spectrum showed positive peaks (red) corresponding with single proton CH correlations at δ 6.86;116.26 (H-3;C-3), δ 6.82;116.08 (H-6;C-6) and δ 6.72;120.46 (H-4;C-4) and negative peaks (blue) corresponding to $CH_2$ correlations at δ 2.82;26.63 (H-7;C-7) and δ 2.64;35.80 (H-8;C-8). The 2- and 3-bond $^{1}$H-$^{13}$C correlations in the 2D $^{1}$H-$^{13}$C HMBC spectrum (green) are marked, allowing the build-up of the compound, fitting the structure of 3-(3,4-dihydroxyphenyl)propionic acid (DHPPA). (**B**) The identity of DHPPA as assigned by 1D and 2D NMR spectroscopy was further confirmed by comparison of the 1D $^{1}$H NMR spectra of the commercially available standard of DHPPA in blue, with the isolated product in red. (**C**) LC-ESI-MS in negative mode showing the exact same mass and retention time as the commercially available standard of DHPPA.

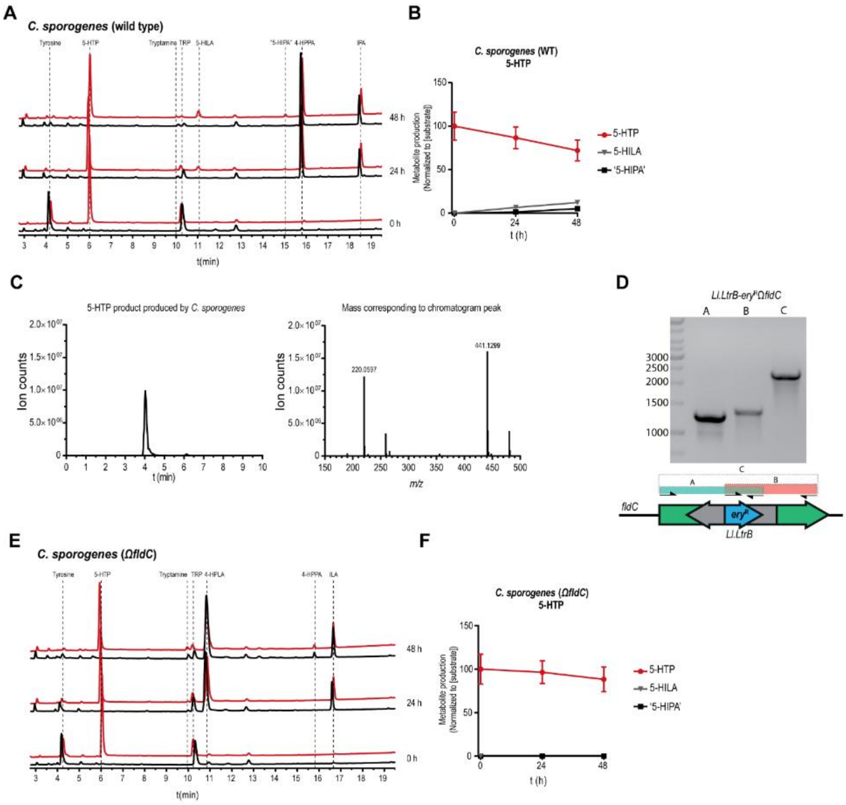

**Supplementary Figure 2. 5-HTP conversion by *Clostridium sporogenes*.** (**A and B**) HPLC-ED curves and quantification (n=3) from supernatant of a *C. sporogenes* batch culture conversion of 5-HTP (5-hydroxy-L-tryptophan) overtime. At the beginning of growth (timepoint 0 h), 100 µM of 5-HTP (red) was added to the culture medium, the black line in the chromatogram depicts the control samples. In 24 h, 5-HTP was converted, to a minor extent, to 5-HILA (5-hydroxyindole-3-lactic acid), as determined by LC-MS and potentially to 5-HIPA (5-hydroxyindole-3-propionic acid), as this peak was absent in $CS^{\Omega fldC}$ incubations. (**B**) Quantification (n=3) of 5-HTP conversion by *C. sporogenes* wild type (also see **Supplementary Table 1**). (**C**) LC-ESI-MS analysis shows the mass of the first peak produced by $CS^{WT}$ from 5-HTP isolated from the HPLC-ED corresponding to the mass of 5-HILA (predicted exact mass 221.069-$[H^+]$). (**D**) Primers targeting the erythromycin cassette in *Ll.LtrB* intron and primers binding outside the cassette were used to confirm the disruption of the *fldC*. (**E**) HPLC-ED chromatograms of $CS^{\Omega fldC}$ incubation with 100 µM of 5-HTP (red) or control (black); no 5-HIPA is detected and tryptophan and tyrosine are converted to their intermediates ILA (indole-3-lactic acid) and 4-HPLA (3-(4-hydroxyphenyl)lactic acid), respectively. The detection of 5-HILA is hampered by the coeluting 4-HPLA. (**F**) Quantification (n=3) of 5-HTP conversion to by *C. sporogenes* Ω*fldC* (also see

**Supplementary Table 1**). (**A, B, E** and **F**)  All experiments were performed in 3 independent biological replicates and means with error bars representing the SEM are depicted.

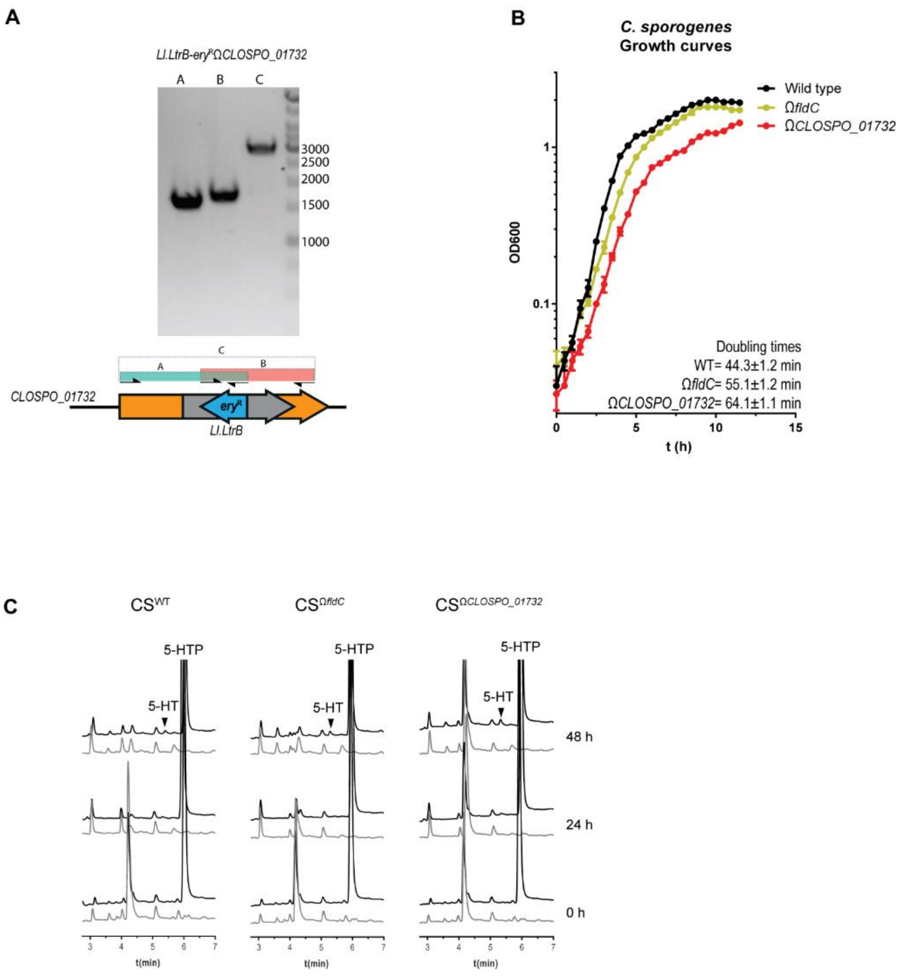

**Supplementary Figure 3. Growth curves of CS$^{\Omega \mathit{fldC}}$ and CS$^{\Omega \mathit{CLOSPO_01732}}$, and 5-HT production.** (**A**) Primers targeting the erythromycin cassette in *Ll.LtrB* intron and primers binding outside the cassette were used to confirm the disruption of the *fldC* and *CLOSPO_01732*. (**B**) Growth-curves of CS$^{WT}$, CS$^{\Omega \mathit{fldC}}$ and CS$^{\Omega \mathit{CLOSPO_01732}}$ showing minor but significant increase in doubling time in the first part of the growth curve. However, all strains reached stationary phase within 12 h. Experiment was performed in triplicate and points and error bars represent the mean with SD (**C**) A minor production of 5-HT (serotonin) is observed in all strains after 48 h, this graph represents 3 independent replicates, see **Supplementary Table 1** for comparison between CS$^{\Omega \mathit{fldC}}$ and CS$^{\Omega \mathit{CLOSPO_01732}}$ with wild type.

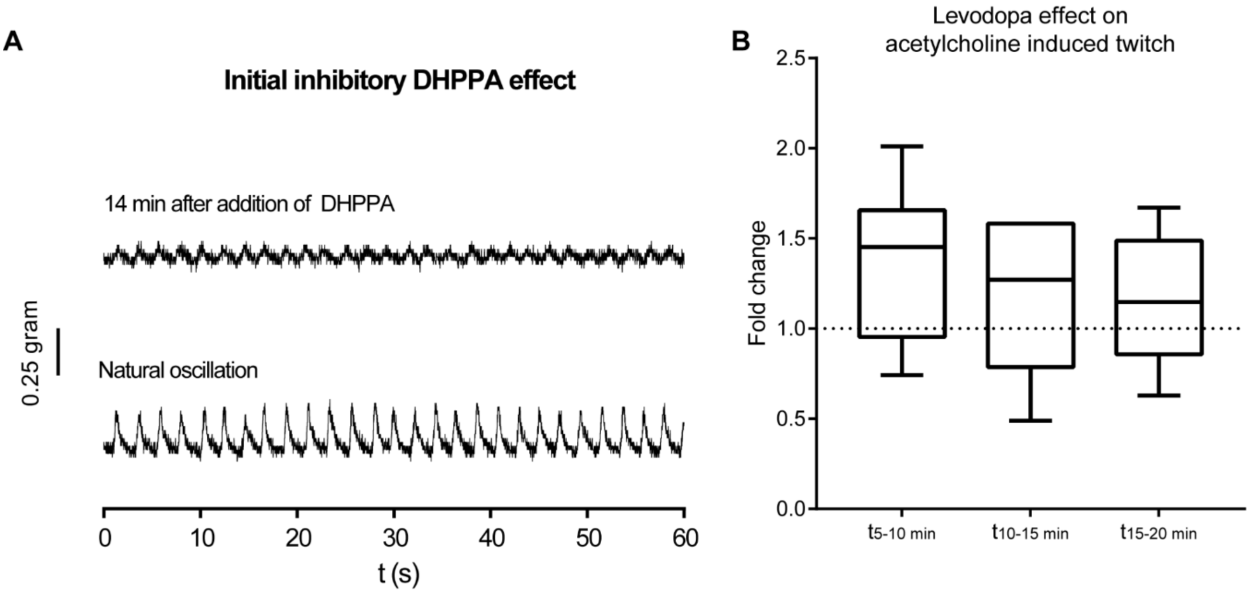

**Supplementary Figure 4. Initial effect of DHPPA on natural ileal contractility and no effect of levodopa on acetylcholine induced twitch.** **(A)** During a period of natural oscillations of contracting ileum 100 µM of DHPPA was added. The amplitude of the contractions decreased and the trace from 14-15 minutes after DHPPA addition is depicted. **(B)** Levodopa has no significant effect on the acetylcholine induced twitch binned in intervals of 5 minutes (n=3 biological replicates and experiments were repeated 2 times per tissue). Significance was tested using repeated measures (RM) 1-way-ANOVA followed by a Tukey's test. Box represents the median with interquartile range and whiskers represent the maxima and minima.

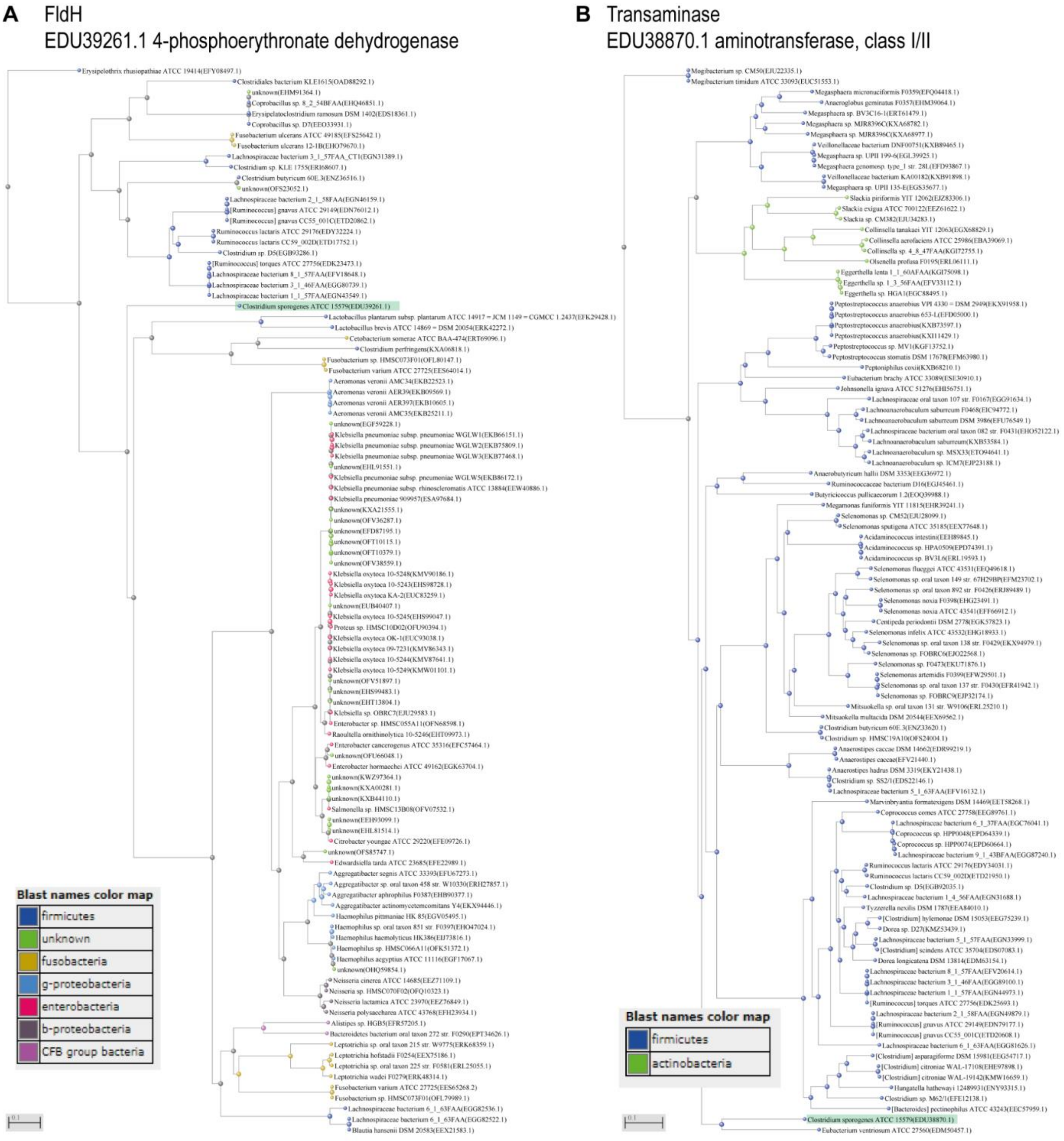

**Supplementary Figure 5. Phylogenetic tree of *C. sporogenes* FldH and EDU38870.** Proteins were BLASTed against the protein sequences from the NIH Human Microbiome Project (HMP) Road map project (PRJNA43021). The top 100 BLASTp hits were aligned in the Constraint-based Multiple Alignment Tool (COBALT) and converted to a distance tree using NCBI TreeView (Parameters: Fast Minimum Evolution; Max Seq Difference, 0.85; Distance, Grishin). In (**A** and **B**) a phylogenetic tree of the top 100 BLASTp hits are depicted for FldH and EDU38870 with *C. sporogenes* indicated by the green bar.

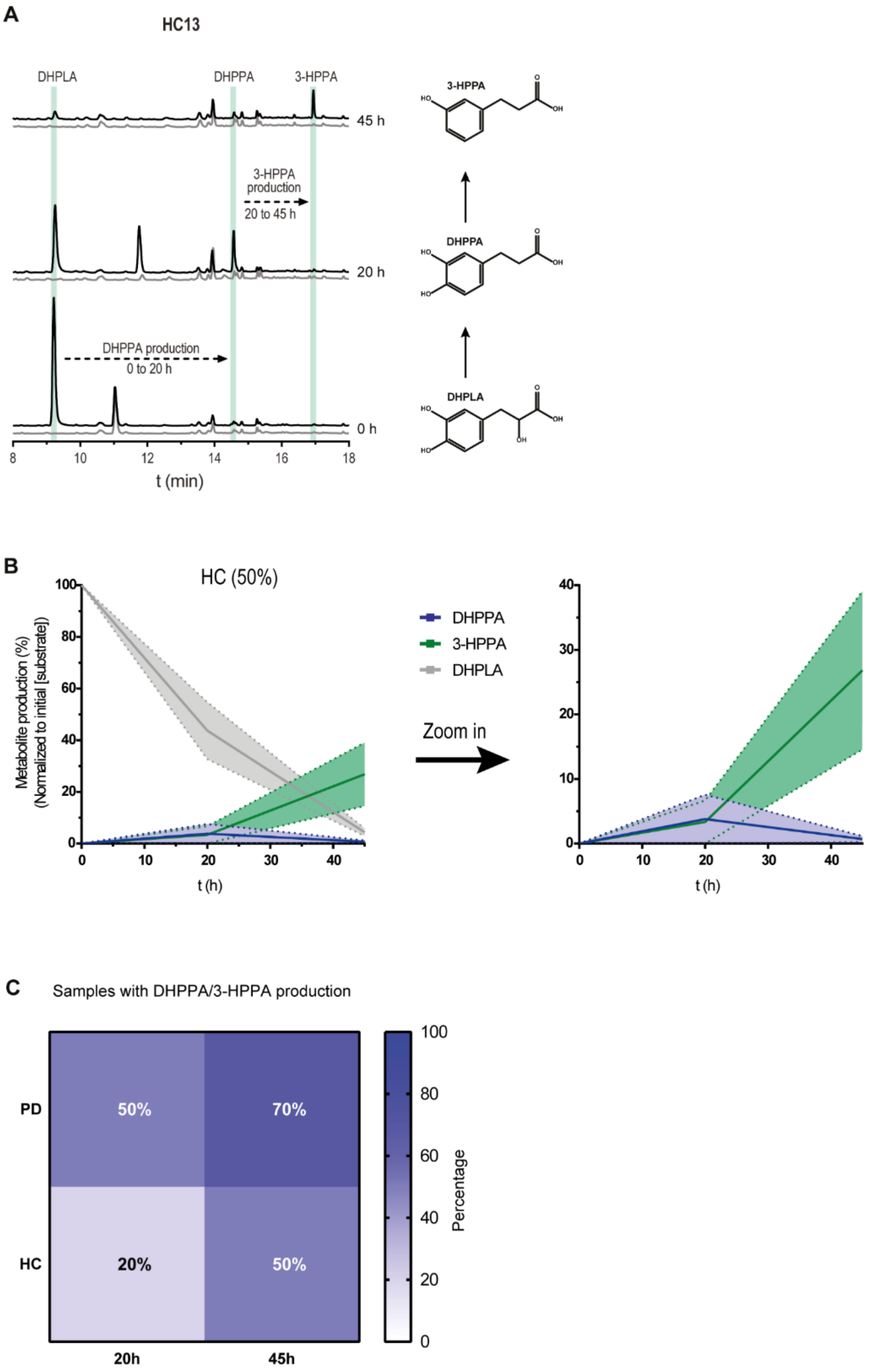

**Supplementary Figure 6. Fecal-incubations from healthy age-matched controls.** (**A**) A representative HPLC-ED chromatogram of fecal-suspension from HC13 where DHPPA is produced from DHPLA (black) after 20 h and is further metabolized to 3-HPPA after 45 h of incubation. The control, without the addition of DHPLA is indicated in grey. Green bars indicate the retention time of the standards indicated. (**B**) Metabolite profiles of the HC fecal suspensions that produced DHPPA or 3-HPPA within 20-45h (50%) are merged as replicates. Lines represent the mean and the shadings the

SEM, a zoom in graph of DHPPA and 3-HPPA is depicted on the right. (**C**) DHPPA or 3-HPPA was quantified as measure for active deamination pathway in the fecal-suspensions. DHPPA or 3-HPPA is produced in 50% and 70% of the PD patient's fecal-suspensions and in 20% and 50% of the HC's fecal-suspensions in 20 and 45 h respectively.

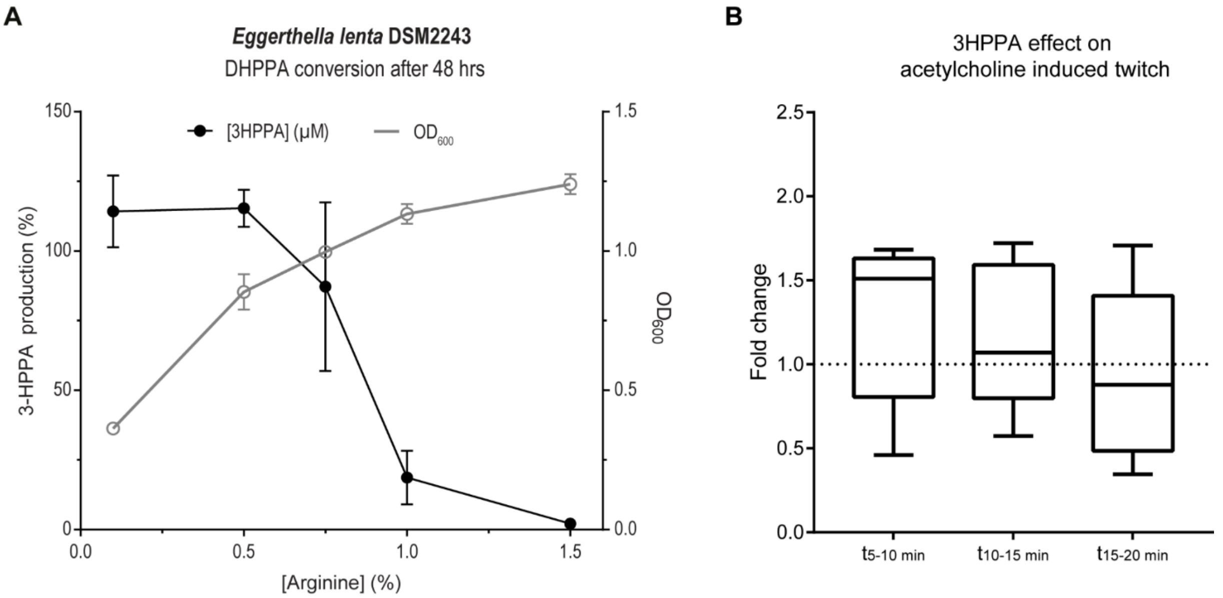

**Supplementary Figure 7. 3-HPPA is produced by *E. lenta*** (**A**) *Eggerthella lenta* has been shown to be able to perform *p*-dehydroxylation of the catecholic B-ring from (+)-catechin and (-)-epicatechin (Jin and Hattori, 2012), a moiety resembling DHPPA. To test whether *E. lenta* could produce 3-HPPA from the dehydroxylation of DHPPA, *E. lenta* DSM 2243 was grown at various concentrations of arginine, which was previously shown to improve growth densities (Haiser et al., 2013). The dehydroxylation of DHPPA to 3-HPPA by *Eggerthella lenta* DSM2243, which is dependent on the arginine concentration in the medium, is shown. The left y-axis indicates 3-HPPA production normalized to initial substrate (DHPPA) concentration. The right y-axis indicates the optical density (OD) at 600 nm. The x-axis indicates the increasing arginine concentration supplied to the medium before 48 h of incubation with 50 µM of DHPPA. At low arginine concentrations, *E. lenta* DSM2243 was capable of dehydroxylation of DHPPA to 3-HPPA, which was inhibited at higher arginine concentrations. Graph represents 3 independent biological replicates and mean with error bars representing the SEM are depicted. (**B**) 3-HPPA has no significant effect on the acetylcholine induced twitch binned in intervals of 5 minutes (n=4 biological replicates and experiments were repeated 2 times per tissue). Significance was tested using repeated measures (RM) 1-way-ANOVA followed by a Tukey's test. Box represents the median with interquartile range and whiskers represent the maxima and minima.

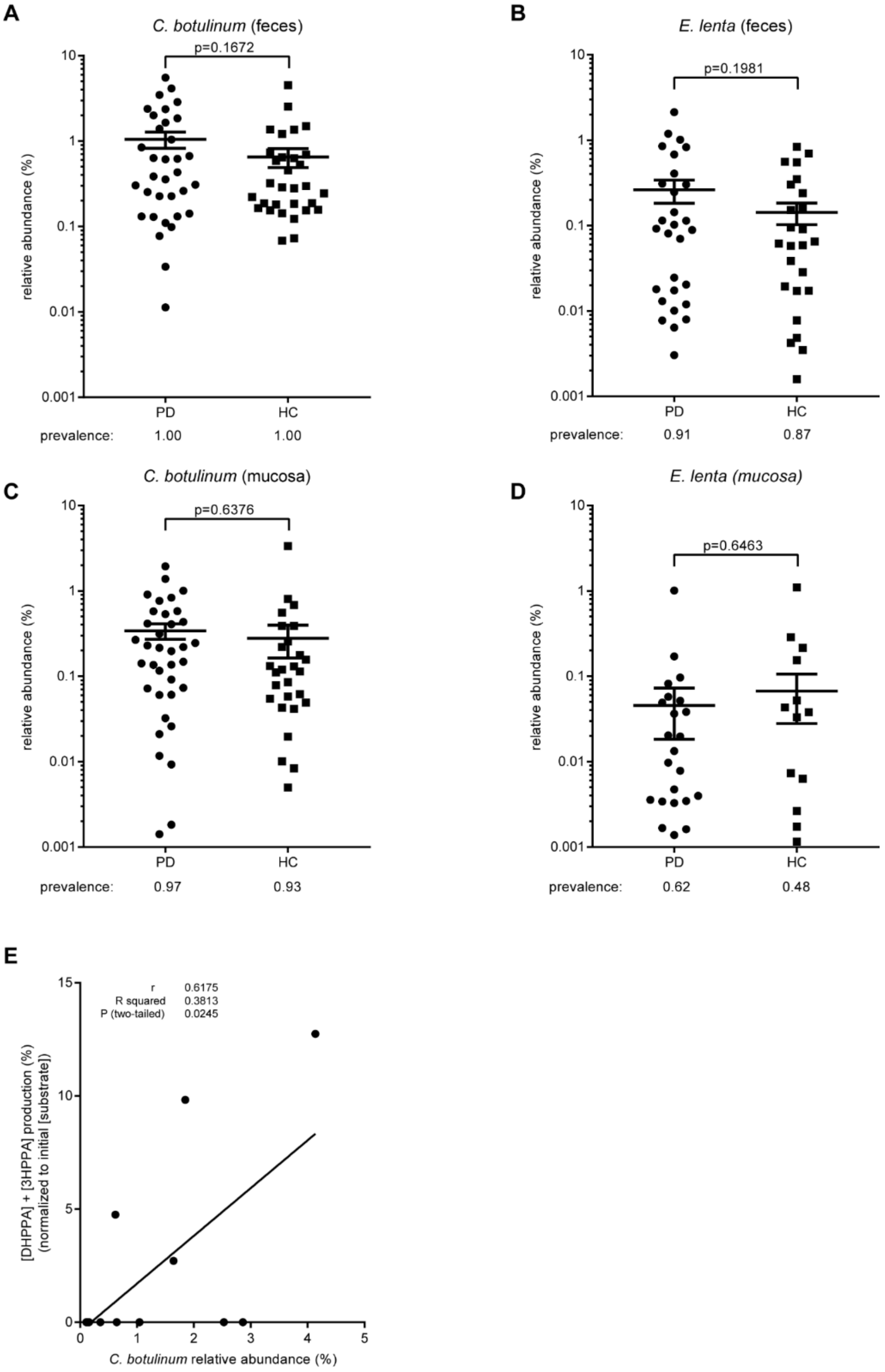

**Supplementary Figure 8. Analysis of 16s rDNA metagenomics data of deaminating bacteria and *E. lenta* in PD and HC fecal and mucosal samples.** (**A**) Relative abundance of *Clostridium botulinum* and (B) *Eggerthella lenta* in PD (feces, n= 34; mucosa, n= 37) and HC (feces, n= 31; mucosa, n=29) fecal samples. (**C**) Relative abundance of *Clostridium botulinum* and (**D**) *Eggerthella lenta* in sigmoid colon mucosa samples from .The cross-header represents the mean and the SEM. (**E**) Linear regression analysis of DHPPA/3HPPA production in PD (n= 9) and HC (n=4) fecal sample incubations at 20 h versus the relative abundance of *C. botulinum* in those samples.

### Supplementary Tables

**Supplementary Table 1.** Values and statistical results corresponding to Figure 2B

| | WT | | | *ΩfldC* | | | ΩCLOSPO_01732 | | |
|---|---|---|---|---|---|---|---|---|---|
| | **0 h** | **24 h** | **48 h** | **0 h** | **24 h** | **48 h** | **0 h** | **24 h** | **48 h** |
| **Phenylalanine** | 100.0 ± 17.7 | n.d. | n.d. | 100.0 ± 15.9 | n.d. | n.d. | 100.0 ± 4.6 | n.d. | n.d. |
| **PLA** | n.d. | n.d. | n.d. | n.d. | 84.7 ± 13.2* | 48.4 ± 9.1* | n.d. | n.d. | n.d. |
| **PPA** | n.d. | 77.6 ± 3.3 | 85.6 ± 0.7 | n.d. | 1.5 ± 0.3**** | 30.7 ± 4.1*** | n.d. | 64.9 ± 3.8# | 71.6 ± 3.1* |
| **Tyrosine** | 100.0 ± 23.0 | 0.4 ± 0.2 | 1.7 ± 1.0 | 100.0 ± 16.8 | 25.1 ± 12# | 4 ± 0.7 | 100.0 ± 18.1 | 110.9 ± 15.9* | 109.3 ± 18.2* |
| **4-HPLA** | n.d. | n.d. | n.d. | 3.2 ± 0.7* | 106.9 ± 15.3* | 113.4 ± 15.0** | n.d. | n.d. | n.d. |
| **4-HPPA** | 1.5 ± 0.3 | 43.7 ± 5.0 | 45.6 ± 4.9 | n.d.* | n.d.** | 2.3 ± 1.1** | n.d.* | 9.2 ± 0.6* | 10.2 ± 0.4** |
| **Tryptophan** | 100.0 ± 21.4 | 19.6 ± 10.0 | 5.4 ± 4.1 | 100.0 ± 20.3 | 47.3 ± 15.3 | 28.3 ± 9.3# | 100.0 ± 16.6 | 70.7 ± 17.5# | 50.9 ± 13.6* |
| **Tryptamine** | n.d. | 6.7 ± 2.3 | 7.0 ± 1.9 | n.d. | 9.6 ± 2.4 | 23.5 ± 2.6* | n.d. | 27.8 ± 1.2** | 40.8 ± 4.4** |
| **ILA** | n.d. | n.d. | n.d. | n.d. | 27.2 ± 1.9*** | 27.0 ± 2.9** | n.d. | n.d. | n.d. |
| **IPA** | 0.6 ± 0.2 | 21.7 ± 1.3 | 22.2 ± 1.8 | n.d.* | n.d.**** | n.d.** | n.d.* | 0.6 ± 0.03**** | 0.8 ± 0.1** |
| **Levodopa** | 100.0 ± 36.4 | 1.0 ± 0.5 | 0.9 ± 0.4 | 100.0 ± 32.7 | 3.4 ± 1.1# | 2.9 ± 0.9# | 100.0 ± 23.1 | 67.5 ± 10.4* | 50.3 ± 8.5* |
| **DHPLA** | n.d. | n.d. | n.d. | n.d. | 172.1 ± 45.2* | 158.6 ± 38.8* | n.d. | n.d. | n.d. |
| **DHPPA** | n.d. | 89.2 ± 24.1 | 84.4 ± 23.1 | n.d. | n.d.* | n.d.* | n.d. | 2.1 ± 0.1* | 2.6 ± 0.6* |
| **5-HTP** | 100.0 ± 16.2 | 86.7 ± 12.6 | 72.1 ± 12.0 | 100.0 ± 17.1 | 96.4 ± 13 | 88.3 ± 14.2 | 100.0 ± 15.4 | 79.7 ± 30.9 | 91.0 ± 14.8 |
| **Serotonin** | 0.04 ± 0.04 | 0.21 ± 0.03 | 0.47 ± 0.07 | 0.04 ± 0.04 | 0.15 ± 0.05 | 0.7 ± 0.11 | 0.06 ± 0.03 | 0.23 ± 0.06 | 1.07 ± 0.12* |
| **5-HILA** | n.d. | 6.6 ± 0.4 | 12.1 ± 0.5 | n.d. | n.d.*** | n.d.**** | n.d. | n.d.*** | n.d.**** |
| **"5-HIPA"** | n.d. | 1.3 ± 0.3 | 5.2 ± 1.3 | n.d. | n.d.* | n.d.* | n.d. | n.d.* | n.d.* |

n.d., not detected; Deamination products are normalized to their initial substrate concentrations (100%). ± values indicate SEM (n=3). Significance was tested between WT and *ΩfldC* or ΩCLOSPO_01732 using a Two-sample equal variance (homoscedastic) Student's t-Test (Microsoft Excel 2019 version 1808). *=p<0.05, **=p<0.0021, ***=p<0.0002, ****=p<0.0001, #=p<0.1234 (not significant).

PLA, 3-phenyllactic acid; PPA, 3-phenylpropionic acid; 4-HPLA, 3-(4-hydroxyphenyl)lactic acid; 4-HPPA, 3-(4-hydroxyphenyl)propionic acid;
ILA, 3-indolelactic acid; IPA, 3-indolepropionic acid; DHLA 3-(3,4-dihydroxyphenyl)lactic acid; DHPPA, 3-(3,4-dihydroxyphenyl)propionic acid;
5-HTP, 5-hydroxytryptophan; 5-HILA, 5-hydroxyindole-3-lactic acid; 5-HIPA, 5-hydroxyindole-3-propionic acid;

**Supplementary Table 2.** MS confirms that DHPPA is extracted from PD and HC samples using alumina extraction method.

| Sample | M-H | ppm error | DHPPA quantification alumina extraction (µM) |
|---|---|---|---|
| DHPPA | 181.0507 | 5.1 | |
| P1 | 181.0509 | 7.5 | 118.2 |
| P2 | 181.0504 | 4.7 | 4.9 |
| P3 | 181.0504 | 4.7 | 1.6 |
| P4 | 181.0505 | 5.3 | 3.8 |
| P5 | 181.0503 | 4.2 | 2.3 |
| P6 | 181.0509 | 7.3 | 154.3 |
| P7 | ND | | 1.4 |
| P8 | 181.0506 | 5.8 | 8.2 |
| P9 | 181.0505 | 5.3 | 11.1 |
| P10 | 181.0504 | 4.7 | 2.4 |
| HC11 | 181.0506 | 5.8 | 2.0 |
| HC12 | 181.0504 | 4.7 | 1.4 |
| HC13 | ND | | 0.5 |
| HC14 | 181.0506 | 5.8 | 1.0 |
| HC15 | ND | | 0.2 |
| HC16 | ND | | 3.5 |
| HC17 | 181.508 | 6.9 | 1.4 |
| HC18 | ND | | 0.5 |
| HC19 | 181.0507 | 5.1 | 4.5 |
| HC20 | 181.0509 | 7.3 | 123.5 |

**Supplementary Table 3 Plasmids and primers used in this study**

| **Plasmid** | **Description** | | **Reference** |
|---|---|---|---|
| pET15b | His-Tag, $amp^R$ | | Novagen |
| pET28b | His-Tag, $kan^R$ | | Novagen |
| pSK023 | pET15b-EDU36436 | | This study |
| pSK024 | pET15b- EDU36793 | | This study |
| pSK025 | pET15b-EDU36848 | | This study |
| pSK026 | pET15b-EDU37030 | | This study |
| pSK027 | pET28b-EDU37032 | | This study |
| pSK028 | pET15b-EDU37374 | | This study |
| pSK029 | pET15b-EDU38761 | | This study |
| pSK030 | pET15b-EDU38870 | | This study |
| pSK031 | pET15b-EDU39385 | | This study |
| **Accession** | **Locus Tag** | **Primers used for cloning (5'-3')** | |
| EDU36436 | CLOSPO_02604 | sk173 FW: GCTACGCATATGAAGTTATCTAAAAAAGCAGTAG<br>sk174 RV: ATTATTCTCGAGCTTTCTAACATTTTATCCACCTC | |
| EDU36793 | CLOSPO_02962 | sk177 FW: GCGCGCCATATGAAAAATAAATTTTTAGCCTATAAG<br>sk178 RV: AATAATCTCGAGCCTCTGAAGCCAAGAAATCTG | |
| EDU36848 | CLOSPO_03017 | sk179 FW: CGCGCGCATATGAAATATGATTTTGATGAAATC<br>sk180 RV: AATAATCTCGAGGCATTTCATAAAAACCCTAGC | |
| EDU37030 | CLOSPO_03199 | sk181 FW: GGCCGCCATATGAAATTTTCAAAAAGAATATCTGACAT<br>sk182 RV: ACGTACCTCGAGGGGTAAGTTCTGAAAATAAAGTA | |
| EDU37032 | CLOSPO_03201 | sk215 FW: CGCGCGCTAGCGTGTTATTTAATGACAAATTAAGAC<br>sk217 RV: GCGCGCCTCGAGTTTATAATATTTATCTAAAACTTTACCTAATC | |
| EDU37374 | CLOSPO_03543 | sk185 FW: GCGCGCCATATGAAGTATAATTTTGACAAAGTAG<br>sk186 RV: GTACACCTCGAGTCCCTCCCATAATTTCAC | |
| EDU38761 | CLOSPO_01623 | sk191 FW: GCAAGCCATATGTTGTTTAAAAAAGGTGGTATTTAT<br>sk192 RV: AAGAATCTCGAGCTTCACTTTAAAGGGAATTTTC | |
| EDU38870 | CLOSPO_01732 | sk193 FW: GCCGGCCATATGATTTCAAATGAAATGCTTAATC<br>sk194 RV: AGTATACTCGAGCAGTTAATTAGCGGTTGTCC | |
| EDU39385 | CLOSPO_00463 | sk197 FW: GCGCATCATATGGATTATATGAAAACTCAAGAAG<br>sk198 RV: AGTAATCTCGAGTCTCAACCTTTAAAGAATGTTAAG | |
| *restrictions sites are underlined | | | |

### Supplementary Methods

#### NMR

Samples were exchanged once with 99.9 atom% $D_2O$ with intermediate lyophilization, finally dissolved in 650 µL $D_2O$. One- and two-dimensional $^{1}$H and $^{13}$C NMR spectra were recorded at a probe temperature of 25°C on a Varian Inova 500 spectrometer (NMR Department, University of Groningen). Chemical shifts are expressed in ppm in reference to external acetone (δ $^{1}$H 2.225; δ $^{13}$C 31.08). 1D 500-MHz 1H NMR spectra were recorded with 5000 Hz spectral width at 16 k complex data points, using a WET1D pulse to suppress the HOD signal. Homonuclear decoupled $^{1}$D 125 MHz $^{13}$C NMR spectra were recorded with 31,000 Hz spectral width at 64k complex data points. 2D $^{1}$H-$^{13}$C HSQC spectroscopy was performed using multiplicity editing, rendering $CH_2$ signals in the negative plane, while CH and $CH_3$ remain in the positive plain. 2D $^{13}$C-$^{1}$H HMBC spectroscopy was performed suppressing single-bond correlations. Spectra were processed using MestReNova v9.1 (Mestrelabs Research SL, Santiago de Compostela, Spain).

### LC-MS

HPLC-MS analysis was performed using an Accella1250 HPLC system coupled with the benchtop ESI-MS Orbitrap Exactive (Thermo Fisher Scientific, San Jose, CA, USA) in negative and positive ion mode. Samples were analyzed on a C18 column (Shim Pack Shimadzu XR-ODS 3 × 75 mm) using a gradient of water/acetonitrile with 0.1% formic acid (0-5 min, 98–90% $H_2O$; 5-10 min, 90-5% $H_2O$; 10-13 min 5% $H_2O$; 13-14 min 98% $H_2O$). Data analysis was performed using Qual Browser Thermo Xcalibur software (version 2.2 SP1.48).

HPLC-MS analysis of alumina extraction samples was performed using an Waters Acquity Class-I UPLC (Waters Chromatography B.V, Etten-Leur, The Netherlands) system coupled to a MaXis Plus Q-TOF (Bruker, Billerica, MA, USA) on negative ion mode with post-column addition of 3 µl/min ESI Tune Mix (G1969-85000; Agilent Technologies, Middelburg, The Netherlands) for mass calibration. Samples were analyzed on a C18 column (Shim Pack Shimadzu XR-ODS 3 × 75 mm) using a gradient of water/acetonitrile with 0.1% formic acid (0-5 min, 98–90% $H_2O$; 5-10 min, 90-5% $H_2O$; 10-13 min

5% $H_2O$; 13-15 min 2% $H_2O$; 15-17 min 98% $H_2O$). Data analysis was performed using Bruker Compass Data Analysis (version 4.2 SR1).

**Bioinformatics**

Phylogenetic trees. Proteins were BLASTed against a local BLAST database constructed from the protein sequences of the NIH Human Microbiome Project (HMP) Roadmap project (PRJNA43021) using BLAST 2.9.0+, NCBI. The top 100 BLASTp hits were aligned in the Constraint-based Multiple Alignment Tool (COBALT, NCBI) and converted to a distance tree using NCBI TreeView (Parameters: Fast Minimum Evolution; Max Seq Difference, 0.85; Distance, Grishin).

Sequence data analysis. The demultiplexed paired-end sequence data from stool and sigmoid colon samples of PD patients and healthy controls from Keshavarzian *et al*., 2015 (bioproject PRJNA268515) were analyzed using Kraken2 (v2.0.9, April 7, 2020), a *k*-mer taxonomic classification system (Wood et al., 2019), using the standard Kraken2-database. To further estimate the species abundance the Kraken2 output was analyzed with Bracken (Bayesian Reestimation of Abundance with KrakEN; v2.6.0, April 3, 2020) (Lu et al., 2017). The number of mapped reads from bacteria with the *fld*-gene cluster (Dodd et al., 2017) were extracted from the Bracken results and the abundance was calculated relative to the total number of mapped bacterial reads.
